# mRNA stem-loops can pause the ribosome by hindering A-site tRNA binding

**DOI:** 10.1101/2020.02.05.936120

**Authors:** Chen Bao, Sarah Loerch, Clarence Ling, Andrei A. Korostelev, Nikolaus Grigorieff, Dmitri N. Ermolenko

## Abstract

Although the elongating ribosome is an efficient helicase, certain mRNA stem-loop structures are known to impede ribosome movement along mRNA and stimulate programmed ribosome frameshifting via mechanisms that are not well understood. Using biochemical and single-molecule Förster resonance energy transfer (smFRET) experiments, we studied how frameshift-inducing stem-loops from *E. coli dnaX* mRNA and the *gag-pol* transcript of Human Immunodeficiency Virus (HIV) perturb translation elongation. We find that upon encountering the ribosome, the stem-loops strongly inhibit A-site tRNA binding and ribosome intersubunit rotation that accompanies translation elongation. Electron cryo-microscopy (cryo-EM) reveals that the HIV stem-loop docks into the A site of the ribosome. Our results suggest that mRNA stem-loops can transiently escape ribosome helicase by binding to the A site. Thus, the stem-loops can modulate gene expression by sterically hindering tRNA binding and inhibiting translation elongation.

## Introduction

During translation elongation, the ribosome moves along mRNA in a codon-by-codon manner while the mRNA is threaded through the mRNA channel of the small ribosomal subunit. In the bacterial ribosome, the mRNA channel accommodates 11 nucleotides downstream of the first (+1) nucleotide of the P-site codon (Takyar et al., 2005; Yusupova et al., 2001). The translating ribosome must unfold mRNA secondary structure to feed single-stranded mRNA through the narrow mRNA channel. The ribosome was shown to be a processive helicase, which unwinds three basepairs per translocation step (Qu et al., 2011; Takyar et al., 2005; Wen et al., 2008). Accordingly, transcriptome-wide ribosome profiling analysis demonstrated that most of secondary structure elements within coding regions of mRNAs do not influence the rate of translation elongation (Del Campo et al., 2015).

Although the elongating ribosome is an efficient helicase, certain mRNA stem-loop structures are known to pause or stall ribosome movement along mRNA. mRNA stem-loop structures can induce ribosome stalling that results in accumulation of truncated polypeptides (Yan et al., 2015) and no-go mRNA decay (Doma and Parker, 2006). In addition, evolutionarily conserved mRNA stem-loops trigger programmed translation pauses. For example, the α subunit of the signal recognition particle receptor is co-translationally targeted to the endoplasmic reticulum membrane by a mechanism that requires a translational pause induced by an mRNA stem-loop structure (Young and Andrews, 1996). Ribosome pausing induced by mRNA structures accompanies −1 programmed ribosome frameshifting (PRF), which controls expression of a number of proteins in bacteria, viruses and eukaryotes (Caliskan et al., 2015). In particular, −1 PRF regulates synthesis of DNA polymerase III in bacteria (Tsuchihashi and Kornberg, 1990); gag-pol proteins of Human Immunodeficiency Virus (HIV) (Jacks et al., 1988), and HIV cytokine receptor ccr5 in higher eukaryotes (Belew et al., 2014).

−1 PRF requires the presence of two signals in an mRNA: the heptanucleotide slippery sequence XXXYYYZ (where X and Z can be any nucleotide and Y is either A or U) and a downstream frameshift stimulating sequence (FSS). The FSS is an RNA hairpin or a pseudoknot (Atkins et al., 2001). The slippery sequence allows cognate pairing of the P-site and A-site tRNAs in both 0 and −1 frames and thus makes frameshifting thermodynamically favorable (Bock et al., 2019). The mechanism by which FSS stimulates frameshifting is less clear. A number of studies have shown that FSSs inhibit the rate of translocation of the A- and P-site tRNAs basepaired with the slippery sequence by at least one order of magnitude (Caliskan et al., 2014; Caliskan et al., 2017; Chen et al., 2014; Kim et al., 2014) and thus produce ribosome pauses (Caliskan et al., 2014; Caliskan et al., 2017; Chen et al., 2014; Kim et al., 2014; Kontos et al., 2001; Lopinski et al., 2000; Somogyi et al., 1993; Tu et al., 1992).

It remains puzzling why certain stem-loops including FSSs induce ribosome pausing in spite of ribosome helicase activity. Slow unwinding of secondary structure, to which ribosome pausing is often attributed, is unlikely to account for the extent of translation inhibition induced by FSSs. Single-molecule experiments showed that translocation through three GC basepairs is only 2 to 3-fold slower than translocation along a single-stranded codon (Chen et al., 2013; Desai et al., 2019; Qu et al., 2011) indicating that the stability of the three basepairs adjacent to the mRNA channel has a relatively moderate effect on translocation rate. Consistent with this idea, neither the thermodynamic stability of the entire FSS nor the stability of the basepairs adjacent to the mRNA entry channel fully correlate with the efficiency of frameshifting (Chen et al., 2009; Hansen et al., 2007; Mouzakis et al., 2013; Ritchie et al., 2012). Hence, FSSs and other stem-loops inducing ribosome pausing may perturb translation elongation by distinct mechanisms that are not well understood.

Here we use a combination of smFRET, biochemical assays and cryo-EM to investigate how FSSs from *E. coli dnaX* mRNA and the *gag-pol* transcript of HIV pause the ribosome. We asked whether these bacterial and viral mRNA sequences, which form stem-loops of the similar lengths, act by a similar mechanism. In agreement with previous studies (Caliskan et al., 2014; Caliskan et al., 2017; Chen et al., 2014; Choi et al., 2020; Kim et al., 2014), we detected FSS-induced inhibition of tRNA/mRNA translocation. We also observed that FSSs inhibit A-site tRNA binding. Cryo-EM analysis of the ribosome bound with FSS-containing mRNA revealed that the FSS docks into the A site of the ribosome and sterically hinders tRNA binding. Occlusion of the ribosomal A site by an mRNA stem-loop may be a common strategy, by which mRNA stem-loops induce ribosome pausing to modulate gene expression.

## Results

### dnaX FSS inhibits intersubunit rotation during translation along the slippery sequence

We investigated how the interaction of FSSs with the ribosome affects cyclic forward and reverse rotations between ribosomal subunits that accompany each translation elongation cycle (Frank and Gonzalez, 2010). Following aminoacyl-tRNA binding to the ribosomal A site and peptide-bond formation, the pre-translocation ribosome predominantly adopts a rotated (R) conformation (Aitken and Puglisi, 2010; Cornish et al., 2008; Ermolenko et al., 2007). In this conformation, the small ribosomal subunit (the 30S subunit in bacteria) is rotated by 7-9° relative to the large subunit (the 50S subunit) (Dunkle et al., 2011; Frank and Agrawal, 2000; Frank and Gonzalez, 2010), and two tRNAs adopt the intermediate hybrid states (Blanchard et al., 2004b; Moazed and Noller, 1989; Valle et al., 2003). EF-G-catalyzed mRNA/tRNA translocation on the small subunit is coupled to the reverse rotation of the ribosomal subunits relative to each other, restoring the nonrotated (NR) conformation in the post-translocation ribosome (Aitken and Puglisi, 2010; Ermolenko et al., 2007; Ermolenko and Noller, 2011).

To probe the effect of FSSs on intersubunit rotation accompanying translation, we employed a model dnaX_Slip mRNA that was derived from the *E. coli dnaX* transcript (Fig. 1). *dnaX* mRNA encodes τ and γ subunits of DNA polymerase III. The γ subunit is produced by a −1 PRF event that occurs with 50-80% efficiency. We chose *dnaX* mRNA because it is one of the most extensively studied −1 PRF systems that has been investigated using both ensemble and single-molecule kinetic approaches (Caliskan et al., 2017; Chen et al., 2014; Choi et al., 2020; Kim et al., 2014; Kim and Tinoco, 2017). The model dnaX_Slip mRNA contained a Shine-Dalgarno (SD, ribosome-binding site) sequence, a short ORF with the slippery sequence AAAAAAG and a downstream 10 basepair-long FSS mRNA hairpin, which together program −1 PRF in *dnaX* mRNA (Larsen et al., 1997) (Fig. 1). In addition, upstream of the SD sequence, the dnaX_Slip mRNA contains a 25 nucleotide-long sequence complementary to a biotin-derivatized DNA oligonucleotide used to tether the mRNA to a microscope slide for smFRET experiments (Fig. 1). The beginning of the ORF encodes Met-Val-Lys-Lys-Arg in 0 frame and Met-Val-Lys-Lys-Glu in −1 frame.

**Figure 1.**
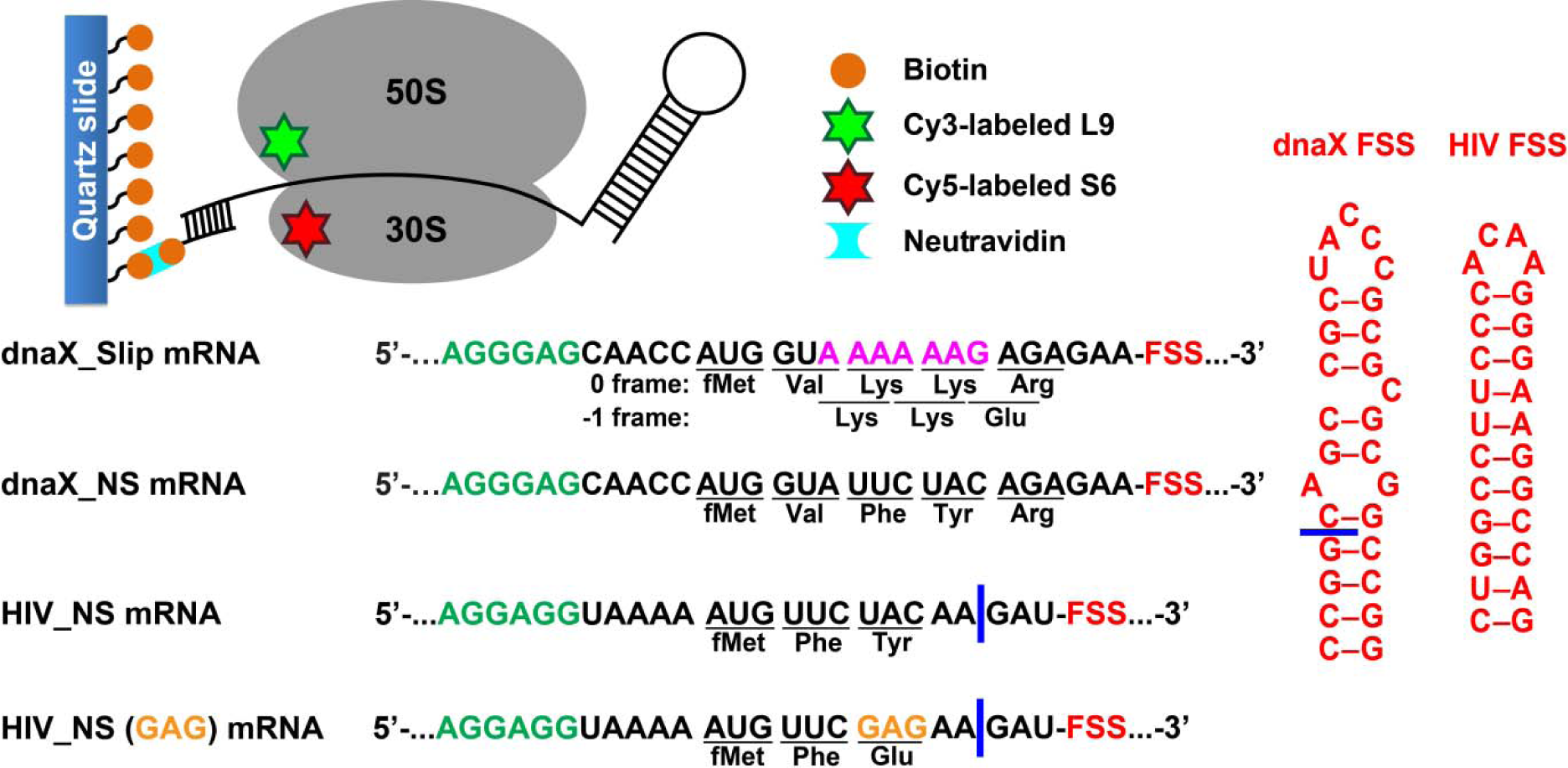
Experimental design. The effect of frameshift-inducing mRNA stem-loops on translation elongation was studied using FRET between cy5 (red) and cy3 (green) attached to 30S protein S6 and 50S protein L9, respectively. S6-cy5/L9-cy3 ribosomes were immobilized on quartz slides using neutravidin and biotinylated DNA oligomers annealed to the mRNA. dnaX_Slip mRNA contains an internal SD sequence (green), a slippery sequence (magenta) and a FSS (red). In the non-slippery (NS) dnaX and HIV mRNAs, the slippery sequences were replaced by non-slippery codons. Two different HIV_NS mRNAs contain either UAC or GAG (orange) codon. Corresponding polypeptide sequences are shown below each mRNA. The ΔFSS mRNAs are truncated as indicated by blue bars.

We determined the efficiency of −1 PRF on the dnaX_Slip mRNA during translation along the slippery sequence via the filter-binding assay. To that end, ribosomes bound with dnaX_Slip mRNA and P-site *N*-Ac-Val-tRNA^Val^ were incubated with EF-G•GTP, EF-Tu•GTP, Lys-tRNA^Lys^, Arg-tRNA^Arg^ (binds in 0 frame) and [^3^H]Glu-tRNA^Glu^ (binds in −1 frame). Consistent with previous publications (Caliskan et al., 2017; Kim and Tinoco, 2017; Larsen et al., 1997), we observed frameshifting efficiency of ∼60% (Suppl. Fig. S1). When ribosomes were programmed with the truncated dnaX_Slip ΔFSS mRNA, which lacks the FSS (Fig. 1), the efficiency of −1 PRF decreased to ∼25%, demonstrating that the FSS stimulates ribosome frameshifting.

To follow intersubunit rotation during translation along the slippery sequence of the dnaX_Slip mRNA, we measured smFRET between fluorophores attached to the 50S protein L9 and the 30S protein S6. The nonrotated (NR) and rotated (R) conformations of the ribosome have been shown to correspond to 0.6 and 0.4 FRET states of S6-cy5/L9-cy3 FRET pair (Cornish et al., 2008; Ermolenko et al., 2007).

We asked whether dnaX FSS positioned near the entrance of the mRNA channel perturbs ribosome intersubunit dynamics during frameshifting. To this end, we monitored elongation on S6-cy5/L9-cy3 ribosomes bound with P-site *N*-Ac-Val-Lys-tRNA^Lys^ and dnaX_Slip mRNA immobilized on a microscope slide (Fig. 1). In this ribosome complex, the second Lys codon of the slippery sequence is positioned in the A site, and the FSS is expected to be one nucleotide downstream of the entrance to the mRNA channel (Yusupova et al., 2001; Zhang et al., 2018). Consistent with previous reports, ribosomes containing P-site peptidyl-tRNA (*N*-Ac-Val-Lys-tRNA^Lys^) are predominately in the NR (0.6 FRET) state (Fig. 2A, Suppl. Fig. S2A) (Cornish et al., 2008; Ermolenko et al., 2007).

**Figure 2.**
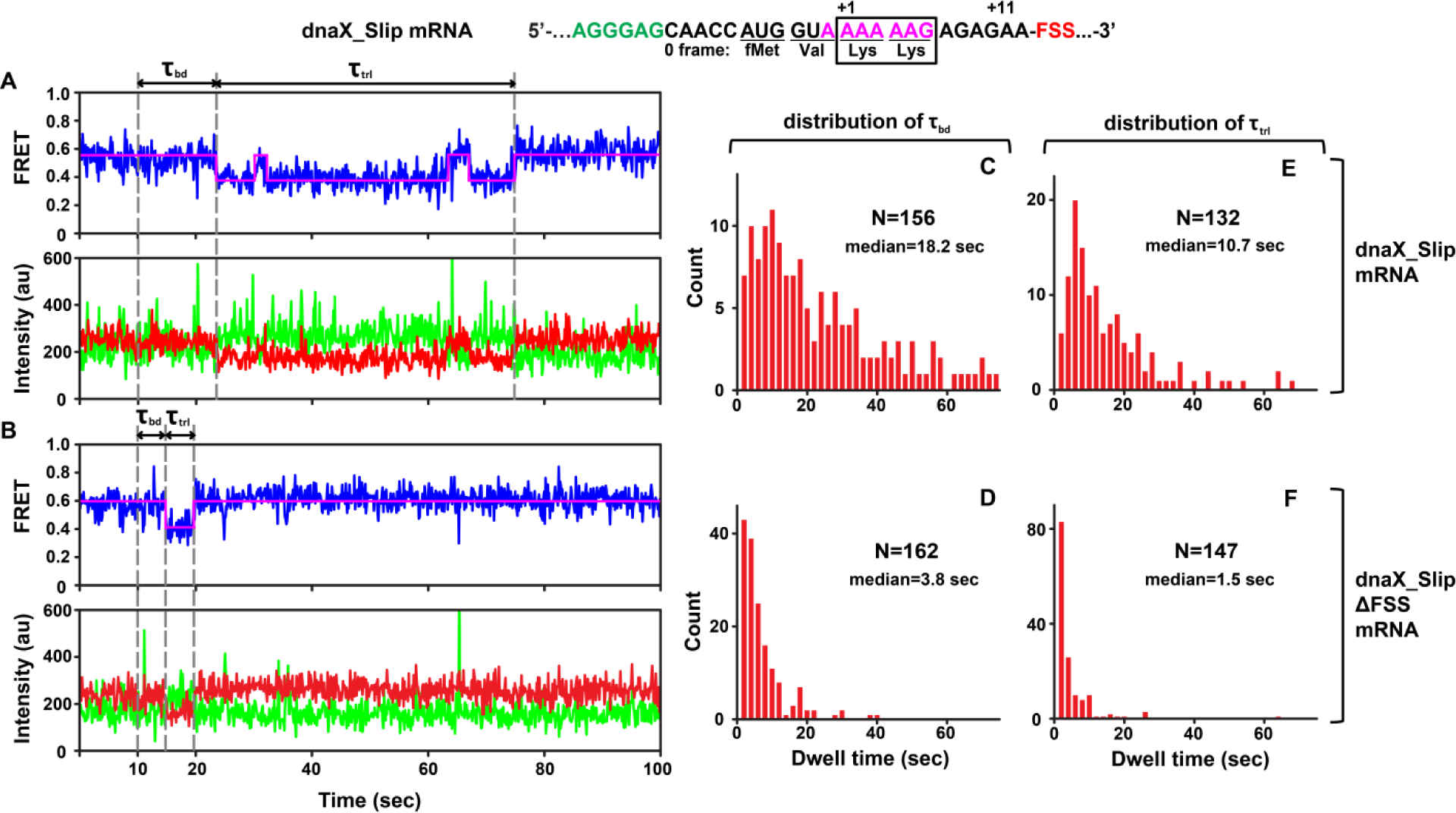
DnaX FSS slows ribosome intersubunit rotation. S6-cy5/L9-cy3 ribosomes containing P-site *N*-Ac-Val-Lys-tRNA^Lys^ were programmed with either dnaX_Slip (A, C, E) or dnaX_Slip ΔFSS (B, D, F) mRNAs. After 10 seconds of imaging, EF-Tu•GTP•Lys-tRNA^Lys^ and EF-G•GTP were co-injected into the flow-through chamber. **(A-B)** Representative smFRET traces show cy3 fluorescence (green), cy5 fluorescence (red), FRET efficiency (blue) and HHM fit of FRET efficiency (magenta). τ_bd_ is dwell time between the injection and Lys-tRNA^Lys^ binding to the A site, which corresponds to the transition from NR (0.6 FRET) to R (0.4 FRET) state of the ribosome. τ_trl_ is dwell time between A-site binding of Lys-tRNA^Lys^ and EF-G-catalyzed tRNA translocation, which corresponds to the transition from R (0.4 FRET) to stable (i.e. lasting over 4 seconds) NR (0.6 FRET) state of the ribosome. The full-length views of smFRET traces are shown in Suppl. Fig. S2 A-B. **(C-F)** Histograms (2-second binning size) compiled from over 100 traces show the distributions and median values of τ_bd_ and τ_trl_. N indicates the number of FRET traces assembled into each histogram.

After 10 seconds of imaging, EF-Tu•GTP•Lys-tRNA^Lys^ and EF-G•GTP were injected to bind Lys-tRNA^Lys^ to the second Lys codon of the slippery sequence and induce tRNA/mRNA translocation. After the injection, the ribosomes showed an NR (0.6 FRET)-to-R (0.4 FRET) transition (Fig. 2A, Suppl. Fig. S2A). The transpeptidation reaction and subsequent movement of tRNAs into hybrid states are typically much faster than tRNA binding to the A site (Blanchard et al., 2004a; Johansson et al., 2008; Juette et al., 2016; Rodnina and Wintermeyer, 2001; Sharma et al., 2016). Hence, the dwell time between the injection and the transition from the NR (0.6 FRET) to R (0.4 FRET) state, τ_bd_, primarily reflects the rate of Lys-tRNA^Lys^ binding to the A site of the ribosome.

The subsequent reverse transition from the R (0.4 FRET) to the stable NR (0.6 FRET lasting over 4 seconds) indicated translocation of mRNA and tRNA. In 64% of traces, the transition from 0.4 FRET to the stable 0.6 FRET state was preceded by one or two short-lived excursions from 0.4 to 0.6 FRET, corresponding to spontaneous fluctuations between the R and NR states, characteristic of pre-translocation ribosomes (Fig. 2A, Suppl. Fig. S2A). This observation is consistent with published smFRET experiments demonstrating that under dnaX FSS-induced pausing, two Lys tRNAs undergo multiple unproductive fluctuations between the hybrid and classical states (Kim et al., 2014).

To further test whether the transition from the R to stable NR conformation indeed corresponds to tRNA translocation, we imaged pre-translocation ribosomes containing deacylated tRNA^Lys^ in the 30S P site in the absence of EF-G. In this complex, R (0.4 FRET) and NR (0.6 FRET) states interconverted at rates of 0.2 sec^-1^ (0.4 FRET to 0.6 FRET) and 0.6 sec^-1^, (0.6 FRET to 0.4 FRET), respectively. Thus, in the absence of EF-G, 95% of pretranslocation ribosomes spent less than 4 seconds in the NR (0.6 FRET) state. This analysis further supports the interpretation that the transition from 0.4 FRET to the stable (i.e. lasting over 4 s) 0.6 FRET state accompanies translocation of tRNAs and mRNA on the small ribosomal subunit (Fig. 2A, Suppl. Fig. S2A). Hence, the dwell time τ_trl_ between the first NR (0.6 FRET) to R (0.4 FRET) transition and the transition from R (0.4 FRET) to the stable NR (0.6 FRET) state in our injection experiments corresponds to the translocation rate.

τ_bd_ and τ_trl_ of dnaX_Slip mRNA programmed ribosomes were remarkably long with median values of 18.2 s and 10.7 s, respectively (Fig. 2C, E), suggesting inefficient tRNA^Lys^ binding and translocation. Moreover, actual median values of τ_bd_ and τ_trl_ are likely longer because Cy5 photobleaching occurring at the rate of 0.02 sec^-1^ leaves some ribosomes with long τ_bd_ and τ_trl_ undetected. Notably, both τ_bd_ and τ_trl_ were broadly distributed (Fig. 2C, E) and could not be fit to a single exponential decay, suggesting heterogeneity within the ribosome population.

Ribosome complexes assembled with dnaX mRNA lacking the FSS (dnaX_Slip ΔFSS mRNA) showed markedly different behavior in comparison with dnaX_Slip mRNA complexes. When EF-Tu•GTP•Lys-tRNA^Lys^ and EF-G•GTP were added to S6-cy5/L9-cy3 ribosomes programmed with dnaX_Slip ΔFSS mRNA and P-site peptidyl-tRNA (*N*-Ac-Val-Lys-tRNA^Lys^), rapid transition from 0.6 to 0.4 FRET was followed by rapid transition to the stable 0.6 FRET (Fig. 2B, Suppl. Fig. S2B). In contrast to dnaX_Slip FSS mRNA programmed ribosomes, which showed spontaneous fluctuations between R and NR states before the transition to the stable post-translocation NR state, only 6% of ribosomes programmed with dnaX_Slip ΔFSS mRNA showed short-lived excursions from 0.4 to 0.6 FRET before translocation. Median values of τ_bd_ (3.8 s) and τ_trl_ (1.5 s) for dnaX_Slip ΔFSS mRNA (Fig. 2D, F) were 5- and 7-fold shorter, respectively, than those measured in ribosomes programmed with dnaX_Slip mRNA. In agreement with previously published results (Caliskan et al., 2017; Chen et al., 2014; Choi et al., 2020; Kim et al., 2014), our data demonstrate that dnaX FSS positioned near the mRNA channel entrance strongly inhibits mRNA/tRNA translocation (Fig. 2E-F). In addition, our data unexpectedly revealed that dnaX FSS also strongly inhibits A-site tRNA binding during the elongation cycle (Fig. 2C-D). Because such FSS-induced inhibition of A-site binding has not been observed before, we further explored this phenomenon using smFRET, biochemical and cryo-EM approaches.

### In the presence of non-slippery sequence, the FSS from *dnaX* mRNA stalls the ribosome in the NR conformation

We next asked whether the spacing between the FSS and the mRNA entry channel of the ribosome affects the ability of the FSS to inhibit A-site binding. Because frameshifting changes the position of the FSS relative to the mRNA entry channel of the ribosome, we aimed to decouple FSS-induced ribosome pausing from frameshifting. To that end, we replaced the two consecutive Lys codons of the dnaX slippery sequence with UUC (Phe) and UAC (Tyr) “non-slippery” codons to create dnaX_NS (“non-slippery”) mRNA (Fig. 1). Mutations in the slippery sequence of *dnaX* were shown to decrease frameshifting efficiency to low (≤ 5%) or undetectable levels (Bock et al., 2019; Caliskan et al., 2017; Kim and Tinoco, 2017; Larsen et al., 1997; Tsuchihashi and Brown, 1992).

When EF-Tu•GTP•Phe-tRNA^Phe^ was added to ribosomes with the FSS positioned four nucleotides away from the entry of the mRNA channel, the ribosome population converted from a predominant NR conformation (0.6 FRET) (Suppl. Fig. S3A, D) to an R (0.4 FRET) conformation (Suppl. Fig. S3B, E) as expected for the “normal” elongation cycle. By contrast, when EF-Tu•GTP•Tyr-tRNA^Tyr^ was added to ribosomes with the FSS positioned one nucleotide away from the entry of mRNA channel, the majority (60%) of ribosomes remained in the NR (0.6) conformation (Fig. 3B). Hence, the encounter of the ribosome mRNA entry channel with the FSS inhibits conversion of the ribosome from the NR to R conformation, which accompanies tRNA binding. Indeed, when EF-Tu•GTP•Tyr-tRNA^Tyr^ was added to ribosomes programmed with dnaX_NS ΔFSS mRNA, which lacks the FSS, the ribosome population was converted from predominately NR (0.6 FRET) (Fig. 3C) to the R (0.4 FRET) conformation (Fig. 3D).

**Figure 3.**
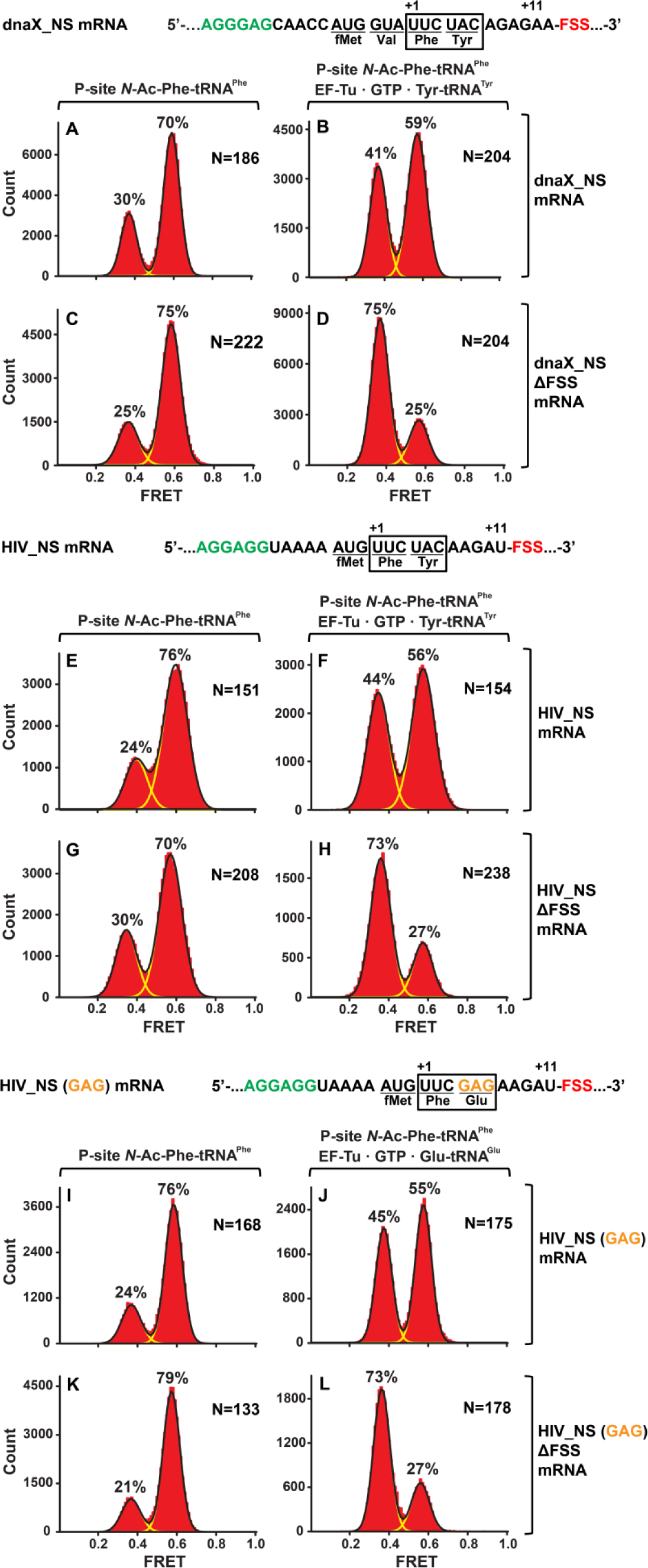
In the context of non-slippery codons, dnaX and HIV FSSs stall the ribosome in NR conformation. Histograms show FRET distributions in S6-cy5/L9-cy3 ribosomes programmed with dnaX_NS **(A-B)**, dnaX_NS ΔFSS **(C-D)**, HIV_NS **(E-F)**, HIV_NS ΔFSS **(G-H)**, HIV_NS (GAG) **(I-J)** or HIV_NS **(GAG)** ΔFSS **(K-L)** mRNA, respectively. Ribosome containing *N*-Ac-Phe-tRNA^Phe^ in the P site **(A, C, E, G, I, K)** were incubated with either EF-Tu•GTP•Tyr-tRNA^Tyr^ **(B, D, F, H)** or EF-Tu•GTP•Glu-tRNA^Glu^ **(J, L)** for 5 minutes and imaged after removal of unbound Tyr-tRNA^Tyr^. Yellow lines show individual Gaussian fits of FRET distributions. Black lines indicate the sum of Gaussian fits. N indicates the number of FRET traces compiled into each histogram. The fractions of the ribosome in R and NR conformations are shown above the corresponding 0.4 and 0.6 Gaussian peaks, respectively.

### The FSS from HIV also stalls the ribosome in the NR conformation

We considered if other frameshift-inducing mRNA stem-loops can induce ribosome stalling in the NR conformation, similar to the FSS from dnaX mRNA. We chose to study the 12 basepair-long RNA hairpin from HIV (Fig. 1) that in combination with the slippery sequence UUUUUUA, induces −1 PRF with 5-10% efficiency to produce the Gag-Pol polyprotein. mRNAs containing the slippery sequence and HIV FSS undergo frameshifting in bacterial (*E. coli*) ribosomes *in vitro* and *in vivo* at frequencies comparable to those observed for HIV frameshifting in eukaryotic translation systems (Brunelle et al., 1999; Korniy et al., 2019; Leger et al., 2004; Mazauric et al., 2009). The FSS from HIV can be studied in *E. coli*, analogous to −1 PRF on mRNA derived from another eukaryotic virus (avian infectious bronchitis virus, IBV) that could also be reconstituted in the *E. coli* translation system (Caliskan et al., 2014) suggesting a common mechanism of frameshifting and ribosomal stalling induced by FSS in bacteria and eukaryotes.

Similar to dnaX_NS mRNA, we designed an HIV_NS mRNA that contained a 25-nucleotide sequence complementary to a biotinylated DNA handle, the SD sequence, and a short ORF containing the FSS. The native slippery sequence, which is positioned 5 nucleotides upstream of the FSS in the HIV mRNA, was replaced with UUC and UAC “non-slippery” codons to delineate the FSS-induced ribosome pausing from frameshifting (Fig. 1). When EF-Tu•GTP•Phe-tRNA^Phe^ was incubated with ribosomes spaced three nucleotides away from the HIV FSS, the conformation of the ribosome population shifted from predominantly NR (0.6 FRET) (Suppl. Fig. S3G, J) to R (0.4 FRET) conformation (Suppl. Fig. S3H, K). By contrast, when EF-Tu•GTP•Tyr-tRNA^Tyr^ was added to ribosomes with the HIV FSS at the entry channel (Fig. 3E), the majority (60%) of ribosomes remained in the NR (0.6 FRET) conformation (Fig. 3F). When EF-Tu•GTP•Tyr-tRNA^Tyr^ was added to ribosomes programmed with HIV_NS ΔFSS mRNA, which lacks the FSS, the ribosome population converted from predominately NR (0.6 FRET) (Fig. 3G) to R conformation (Fig. 3H). Therefore, similar to the FSS from *dnaX*, upon encountering the ribosome, the FSS from HIV stalls the ribosome in the NR conformation.

To test whether identities of A-site codon and A-site tRNA affect the observed ribosome stalling in the NR conformation, we made HIV_NS (GAG) mRNA, in which the original UAC (Tyr) codon of HIV_NS mRNA was replaced with a GAG (Glu) codon (Fig 1). The resulting complexes behaved similarly to the complexes assembled with the original HIV_NS mRNA (Fig. 3I-L). The majority of ribosomes (60%) with the FSS at the mRNA entry channel remained in the NR (0.6 FRET) conformation after addition of EF-Tu•GTP•Glu-tRNA^Glu^ (Fig. 3J) while ribosomes programmed with HIV_NS (GAG) ΔFSS mRNA switched to the R conformation (Fig. 3L). Thus, we show that the FSS-induced inhibition of tRNA binding is independent of A-site codon identity.

Next, we tested whether the stalling in the NR conformation observed with dnaX_NS and HIV_NS mRNAs was due to mRNA frameshifting that prevented A-site binding of Tyr-tRNA^Tyr^ (or Glu-tRNA^Glu^ in the case of ribosomes programmed with HIV_NS (GAG) mRNA). Control experiments described in Supplementary Materials showed that in the absence of the slippery sequence, FSS-induced frameshifting is negligible and does not account for the FSS-induced ribosome stalling in the NR conformation (Suppl. Fig. S4).

HIV and dnaX FSSs placed near the entry to the mRNA channel could stall the ribosome in the NR conformation by either (i) inhibiting A-site tRNA binding, (ii) blocking the peptidyltransfer reaction after the binding of A-site tRNA or (iii) stabilizing the pre-translocation ribosome in the NR conformation. Experiments described in Supplementary Materials demonstrated that HIV and dnaX FSSs placed near the entry to the mRNA channel neither block the peptidyltransfer reaction after the binding of A-site tRNA nor stabilize the pre-translocation ribosome in the NR conformation (Suppl. Fig. S3C, F, I, L and S5A). These results support the idea that HIV and dnaX FSSs stall the ribosome in the NR conformation by inhibiting A-site tRNA binding.

### dnaX and HIV FSS inhibit tRNA binding to the A site of the ribosome

To further test whether FSSs positioned near the entry of the mRNA channel inhibit tRNA binding, we used a filter-binding assay to measure binding of radio-labeled aa-tRNA during translation through four (Met, Val, Phe and Tyr) consecutive codons along the dnaX_NS mRNA. The ribosomes containing P-site *N*-Ac-Met-tRNA^Met^ were then incubated with EF-G•GTP, EF-Tu•GTP, Val-tRNA^Val^, Phe-tRNA^Phe^ and Tyr-tRNA^Tyr^ before loading ribosomes onto a nitrocellulose filter and washing away unbound aa-tRNA. The experiment was repeated three times with one of the three aminoacyl-tRNAs radio labeled, i.e. using [^14^C]Val-tRNA^Val^, [^3^H]Phe-tRNA^Phe^ or [^3^H]Tyr-tRNA^Tyr^. Similar experiments were performed with ribosomes programmed with HIV_NS mRNA to measure binding of radio-labeled aa-tRNA during translation through three (Met, Phe and Tyr) consecutive codons. In ribosomes programmed with dnaX_NS mRNA, binding of [^3^H]Tyr-tRNA^Tyr^ was considerably diminished while the incorporation of Val and Phe into the polypeptide chain were only mildly inhibited compared to levels measured in ribosomes programmed with dnaX_NS ΔFSS mRNA lacking the FSS (Fig. 4A). Likewise, in ribosomes programmed with HIV_NS mRNA, binding of [^3^H]Tyr-tRNA^Tyr^ was strongly inhibited while binding [^3^H]Phe-tRNA^Phe^ was unaffected (Fig. 4B). Hence, consistent with smFRET experiments, the filter-binding assay suggests that both dnaX and HIV FSSs inhibit A-site tRNA binding only when positioned near the entry of the mRNA channel. By contrast, when positioned three or more nucleotides away from the entrance of the mRNA channel at the beginning of elongation cycle, dnaX and HIV FSSs do not perturb A-site binding.

**Figure 4.**
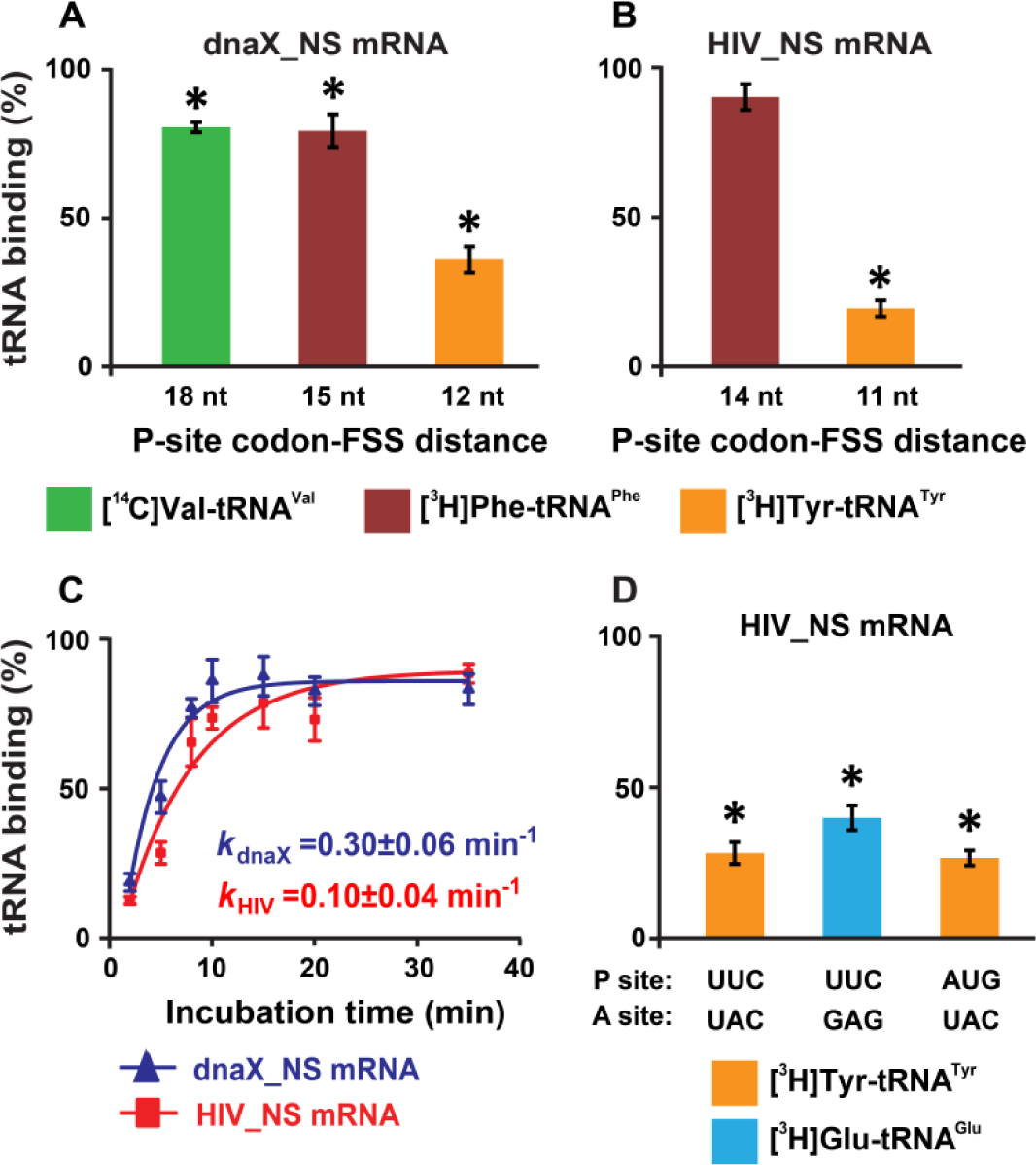
DnaX and HIV FSSs inhibit A-site tRNA binding. **(A-B)** Incorporation of radio-labeled amino acids during translation through first four codons of dnaX_NS mRNA **(A)** or first three codons of HIV_NS mRNA **(B)** were measured by filter-binding assays. **(C)** Kinetics of EF-Tu-catalyzed [^3^H]Tyr-tRNA^Tyr^ binding to the A site of ribosomes containing *N*-Ac-Phe-tRNA^Phe^ in the P site. Ribosomes were programmed with dnaX_NS mRNA (blue) or HIV_NS mRNA (red). Single exponential fits are shown as line graphs. **(D)** The extent of EF-Tu-catalyzed cognate aminoacyl-tRNA binding after 5 minute incubation with ribosomes programmed with HIV_NS, HIV_NS (GAG) or HIV_NS (AUG) mRNAs. The P site of the ribosome was bound with *N*-Ac-Phe-tRNA^Phe^ (in the presence of HIV_NS and HIV_NS (GAG) mRNAs) or *N*-Ac-Met-tRNA^Met^ (in the presence of HIV_NS (AUG) mRNA). (**A-D**) The binding of radio-labeled amino acids to ribosomes programmed with FSS-containing mRNA is shown relative to that observed in ribosomes programmed with corresponding ΔFSS mRNA. Asterisks indicate that amino acid incorporation into ribosomes programmed with FSS-containing mRNA was significantly different from that in ribosomes programmed with ΔFSS mRNA, as *p*-values determined by the Student t-test were below 0.05. Error bars in each panel show standard deviations of triplicated measurements.

Next, we examined the kinetics of [^3^H]Tyr-tRNA^Tyr^ binding to the A site of ribosomes, which contained P-site *N*-Ac-Phe-tRNA^Phe^ and were programmed with either dnaX_NS or HIV_NS mRNA. Both dnaX and HIV FSSs dramatically slowed the rate of [^3^H]Tyr-tRNA^Tyr^ binding down as the apparent pseudo first order rate of tRNA binding was reduced to 0.3 and 0.1 min^-1^, respectively (Fig. 4C). When ribosomes were programmed with either dnaX_NS ΔFSS or HIV_NS ΔFSS mRNAs, the rate of [^3^H]Tyr-tRNA^Tyr^ binding was too fast to be measured by filter-binding assay, which involves manual mixing of ribosomes and tRNA. These kinetic experiments also show that while dnaX and HIV FSSs strongly inhibit A-site tRNA binding, they do not completely block it.

Furthermore, we found that in the absence of EF-Tu, FSSs from dnaX and HIV also inhibit non-enzymatic binding of *N*-Ac-[^3^H]Tyr-tRNA^Tyr^ to the A site of ribosomes programmed with dnaX_NS mRNA or HIV_NS mRNA (Suppl. Fig. S5B). Therefore, the FSS-induced inhibition of A-site tRNA binding is independent of the EF-Tu function. In addition, we observed that FSS-induced inhibition of cognate aminoacyl-tRNA binding to the A site is largely independent of identities of either A- or P-site codons (Fig. 4D). FSS-induced binding inhibition of cognate aminoacyl-tRNA to the A site was observed when filter-binding experiments were performed with ribosomes programmed with mRNAs in which the original UAC (Tyr) or UUC (Phe) codons of HIV_NS mRNA were replaced with GAG (Glu) codon or AUG (Met) codon, respectively (Suppl. Table 1).

Taken together, our smFRET and filter-binding experiments indicate that when positioned 11-12 nucleotides downstream of the first nucleotide of the P-site codon, the FSSs from HIV and *dnaX* mRNAs can substantially inhibit binding of aminoacyl-tRNA to the A site of the ribosome. Assuming that *dnaX* and HIV mRNA are threaded through the 30S mRNA channel, an 11-12 nucleotide distance from the P-site codon corresponds to positioning of the FSSs at the entry of the mRNA channel. Consistent with this hypothesis, a recent cryo-EM reconstruction revealed that the *dnaX* FSS placed 12 nucleotides downstream from the P site codon interacts with ribosomal proteins uS3, uS4, and uS5 located at the 30S mRNA entry channel (Zhang et al., 2018). However, the mRNA entry channel is ∼20 Å away from the 30S decoding center. How the FSS positioned at the mRNA entry channel inhibits tRNA binding to the A site remains unclear.

### Cryo-EM analysis reveals HIV FSS hairpin binding to the A site

To investigate the structural basis for the tRNA binding inhibition by the FSSs, we performed single-particle cryo-EM of the HIV FSS mRNA-ribosome complex. We prepared a 70S *E.coli* ribosome complex programmed with HIV_NS (GAG) mRNA (Fig. 1) and bound with a peptidyl-tRNA analog, *N*-Ac-Phe-tRNA^Phe^, in the P site. Our smFRET and filter-binding experiments showed that in this complex, the HIV FSS inhibits binding of Glu-tRNA^Glu^ to the Glu (GAG) codon in the A site (Fig. 3-4).

Maximum-likelihood classification of a 640,261-particle data set revealed predominant ribosome states that contained strong density in both the P and A sites, which we interpreted as P-site tRNA and the FSS hairpin, respectively (64% particles total). Two classes comprise the ribosome in classical NR states (NR-I and NR-II, ∼1° 30S rotation) with P/P tRNA (Fig. 5A, Suppl. Fig. 6A), while one class represents a R ribosome state with P/E tRNA (R-I, ∼7° 30S rotation) (Fig. 5B, Suppl. Fig. 6B), at overall resolutions between 3.1 Å and 3.4 Å. Additional classes contained weaker density in the A site, likely reflecting compositional and/or conformational heterogeneity (see Methods). By contrast, there is no density at the entry site of the mRNA channel, indicating that the HIV FSS does not bind to the mRNA entry channel.

**Figure 5.**
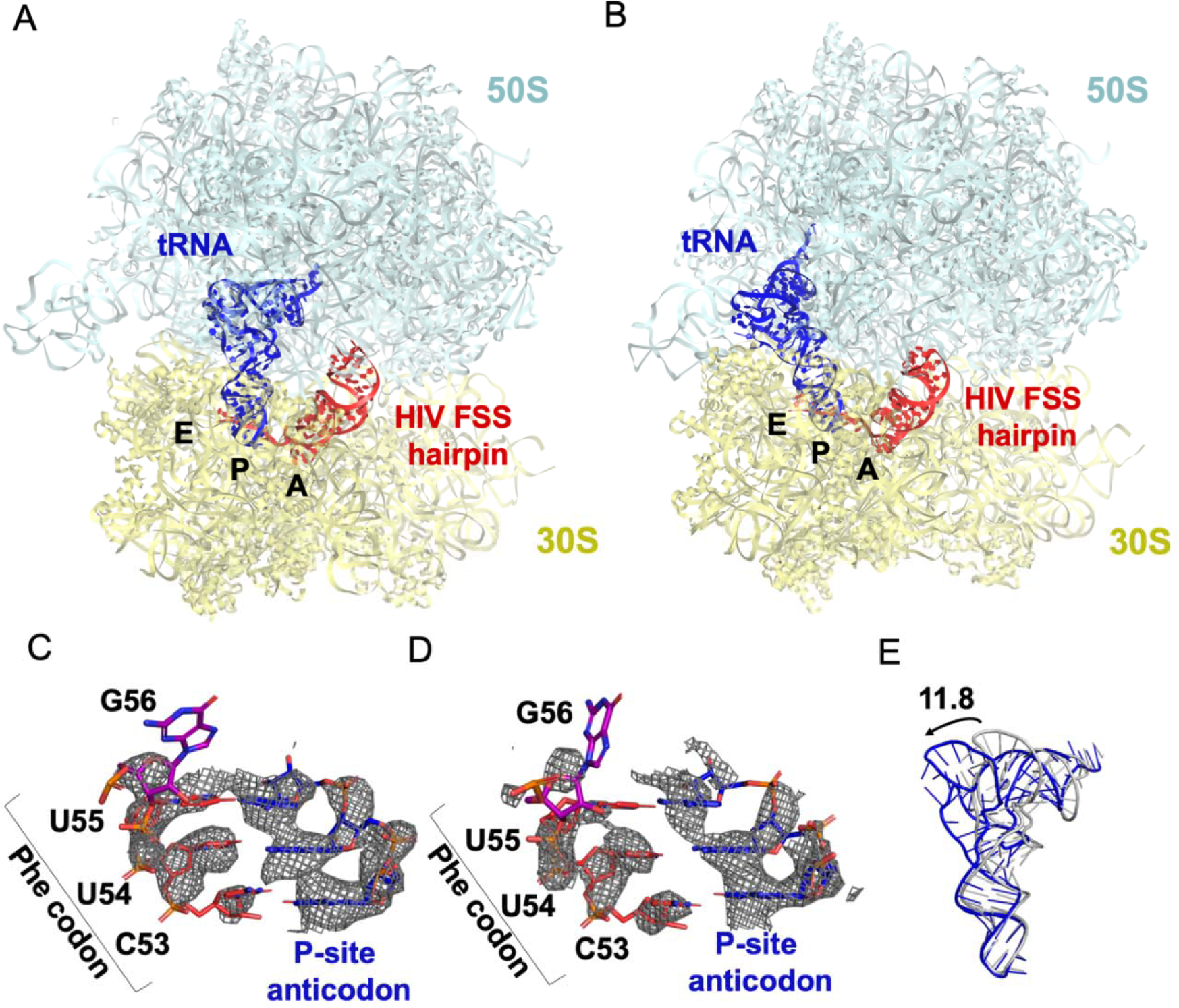
The HIV FSS hairpin occupies the ribosomal A site. **(A)** Cryo-EM structures of the 70S ribosome in non-rotated (NR-I) and **(B)** rotated (R-I) conformations. The large subunits (50S) are shown in aqua, the small subunits (30S) in yellow, and P-site tRNA in blue, HIV FSS hairpin in red. **(C)** and **(D)** Close-up views of the codon and anti-codon basepairs of the NR-I (C) and R-I (D) states illustrating in-frame basepairing off the HIV_NS(GAG) mRNA (red) with the P-site tRNA (blue). The first position of the GAG A-site codon is shown in purple. The cryo-EM map (gray mesh) was sharpened by applying a B-factor of −50 Å^2^. **(E)** Overlay of NR-I P-site tRNA with P-site tRNA bound in the P/P classical site (PDB ID: 4V5D) shows a 14.8 Å rotation of the tRNA elbow towards the E site.

P-site tRNA is base paired with mRNA, indicating the absence of frameshifting. In all three classes NR-I, NR-II and R-I, density allows for the distinction of purines from pyrimidines (Fig. 5C-D), revealing Watson-Crick pairing between *N*-Ac-Phe-tRNA^Phe^ and an in-frame UUC codon. This is consistent with smFRET data showing that in the absence of the slippery sequence, *dnaX* and HIV FSSs do not promote frameshifting (Suppl. Fig. 4).

In both NR-I and NR-II structures, the P-site tRNA elbows are shifted by 11.8 Å and 13.0 Å towards the E site compared to P-site tRNA in other classical states (corresponding to tRNA rotation by 18.5° and 21.0°, respectively compared to PDBID: 4V5D, Fig. 5E). Similar tRNA states were observed in several termination complexes where they are thought to represent an intermediate to the P/E hybrid-state after deacylation by a release factor (Graf et al., 2018; Svidritskiy et al., 2019) (Supp. Fig. 6C). The peptidyl moiety is unresolved in both NR-I and NR-II structures, so it is unclear whether the peptidyl moiety is disordered or hydrolyzed leading to deacylation of the P-site tRNA. Nevertheless, the position of the tRNA CCA tail in the 50S peptidyl-transferase center is similar to that in the peptidyl-tRNA complexes (Polikanov et al., 2014), suggesting that NR-I and NR-II can be sampled with peptidyl-tRNAs. Furthermore, superposition with the P/P-tRNA bound structures demonstrates the absence of steric clash between the hairpin and tRNA, indicating that the hairpin in the A site is also compatible with the classical NR ribosome.

**Figure 6.**
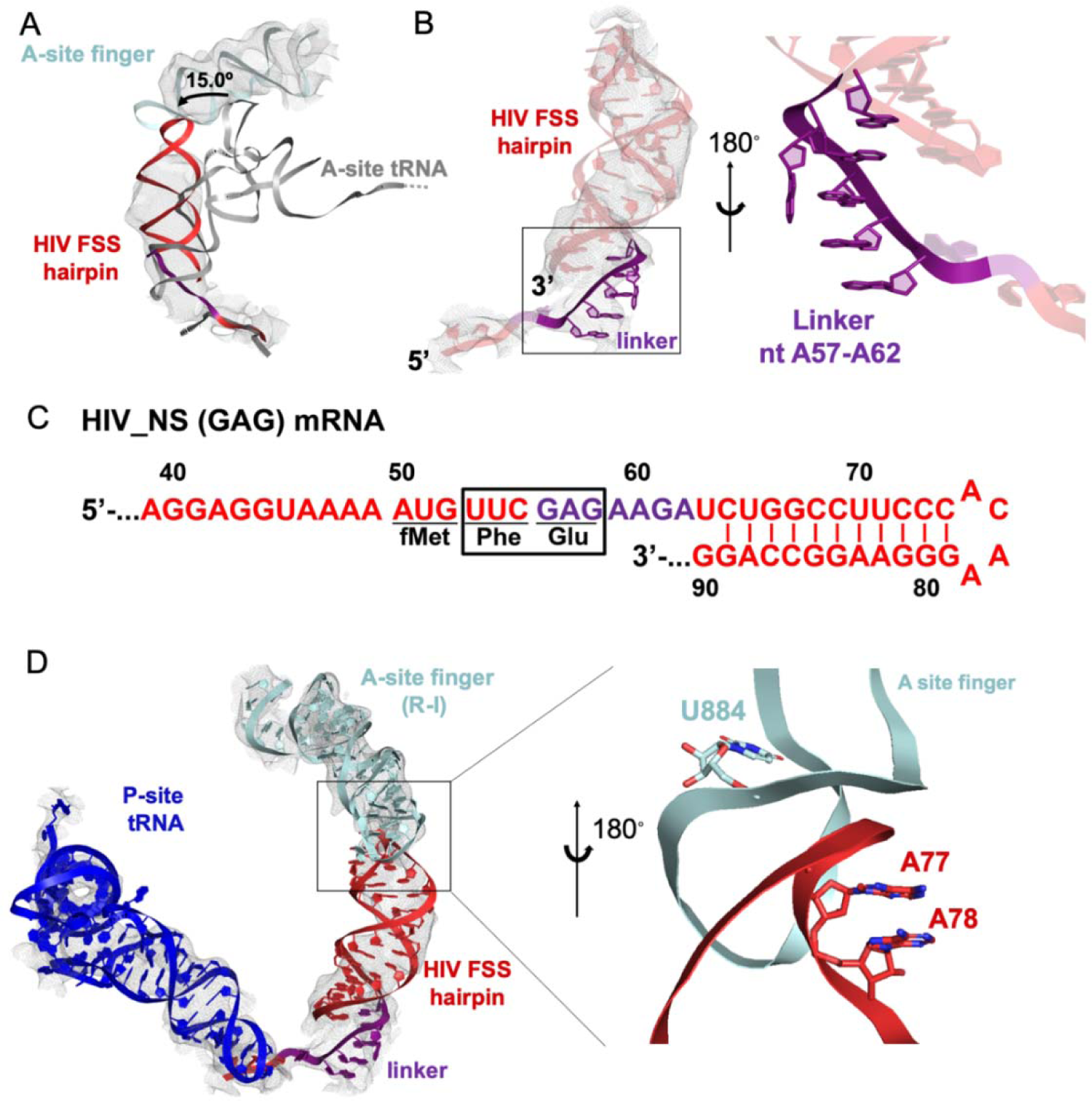
Stacking interactions of linker nucleotides stabilize the HIV FSS in the A site. **(A)** Overlay of the NR-I HIV FSS hairpin from this work with A-site tRNA (grey) accommodated in a ribosome in the classical state (PDB ID: 4V5D). The hairpin density is shown after filtering to 8 Å. **(B)** View of the R-I HIV FSS hairpin model (red, linker in purple) in cryo-EM density filtered to 5 Å (grey mesh) and close-up of the purine stack (shown in purple) after 180° rotation. **(C)** Primary sequence and secondary structure of the HIV_NS(GAG) mRNA. The linker sequence is highlighted in purple. **(D)** In the rotated state, the HIV FSS hairpin (red) contacts the A-site finger (aqua) of the large ribosomal subunit. The hairpin density allows to clearly identify helical pitch, major and minor grooves of the A-form RNA. The close-up after 180° rotation shows that the only complementary bases within the two loop regions point away from each other and likely do not contribute to A-site finger/hairpin binding. P-site tRNA is blue.

Our cryo-EM reconstructions reveal the structural basis for FSS binding to the ribosome. Instead of binding next to the mRNA channel, the FSS hairpin stacks on the purine-rich GAG codon and nucleotides AAGAU (nucleotides 59-63 of HIV (GAG) NS mRNA between GAG and the FSS), which we further refer to as the linker sequence, to occupy the ribosomal A site (Fig 6B-C). To exclude the possibility that the A-site density is an unusually accommodated A-site tRNA, we docked a tRNA into this density. In both the R and NR states, no additional density is observed that could correspond to the acceptor stem of A-site tRNA. Furthermore, extended portions of the tRNA elbow and acceptor stem would clash with the A-site finger of the 50S subunit (Fig 6A, Supp. Fig. 6D-E), because the density is rotated by ∼15° towards the A-site finger (NR-I) compared to A-site tRNA in the classical state (Fig. 6A). In the R-I conformation, the hairpin density merges with the A-site finger (helix38) density suggesting that the hairpin loop (A75-A78) comes into a closer contact with the A-site finger (helix 38) (Fig. 6D) and does not correspond to a hybrid-state A/P tRNA. Thus, our interpretation of the density rules out tRNA in A site and explains inhibition of A-site tRNA binding by the presence of the FSS hairpin.

The local resolution of the A site allows to distinguish the major and minor grooves of the A-form RNA, but individual basepair locations cannot be determined, suggesting that the FSS hairpin samples an ensemble of conformations. We were able to build a plausible pseudoatomic model of the linker and hairpin based on resolved density features and previously solved structures. We used a published NMR model of the HIV FSS hairpin (Staple and Butcher, 2003), which includes the hairpin loop and adjacent nine basepairs (nucleotides 66-87) of the predicted 11 canonical and additional G-U pair (predicted using mfold (Zuker, 2003)). The closing hairpin basepairs, A-site codon (GAG) and the linker sequence were modeled manually to form stacking interactions. The density suggests a dynamic conformation of the first A-site codon position (G56). The following six nucleotides (A57-A62, sequence: AGAAGA) contain an adenine-rich homopurine sequence, which typically adopts stacked A-form-like conformations (Isaksson et al., 2004). Accordingly, we modeled the second and third positions of the A-site codon and the linker as an A form-like purine stack (Fig. 6B). A kink in the density suggests that one nucleotide in the purine stack, possibly A60, is flipped out of the helix. Nucleotides G61 and A62 might form heteropurine basepairs with A92 and G91, respectively, though specific interactions are not visible at this resolution.

Conformational heterogeneity of the hairpin observed in our cryo-EM structures is likely important for the hairpin’s mechanism of action. To allow FSS hairpin entry into the A site, the mRNA linker has to be released from the mRNA channel in the 30S subunit. Dynamic occupancy of the A site by the FSS hairpin may allow an incoming tRNA to eventually overcome the translational block that the hairpin imposes on the ribosome. Binding of tRNA likely stabilizes the A-site codon, allowing the linker to reestablish its position in the mRNA channel and the hairpin to bind next to the channel entry, restoring an elongation ribosome state.

## Discussion

In this study, we investigated molecular mechanisms by which FSSs from *E. coli dnaX* and HIV mRNAs induce ribosome pausing. Although the sequences of these mRNA elements are different, they appear to act via the same mechanism due to similar hairpin structures. We found that when positioned 11-12 nucleotides downstream of the P-site codon, the FSSs perturb translation elongation through two parallel pathways: (i) inhibiting tRNA binding to the A site of the ribosome and (ii) inhibiting ribosome translocation (Fig. 7). These observations support the idea that FSSs stimulate frameshifting by pausing the ribosome. Our findings are consistent with studies demonstrating that −1 PRF on *dnaX* and HIV mRNAs may occur through two pathways: (i) slippage of the single P-site tRNA when the A site remains vacant, or (ii) frameshifting of both P-site and A-site tRNAs when two tRNAs are bound to the ribosome (Caliskan et al., 2017; Korniy et al., 2019). Our finding that the dnaX FSS slows the rate of ribosome translocation by ∼10-fold is also in excellent agreement with a number of previous reports (Caliskan et al., 2017; Chen et al., 2014; Choi et al., 2020; Kim et al., 2014; Kim and Tinoco, 2017).

**Figure 7.**
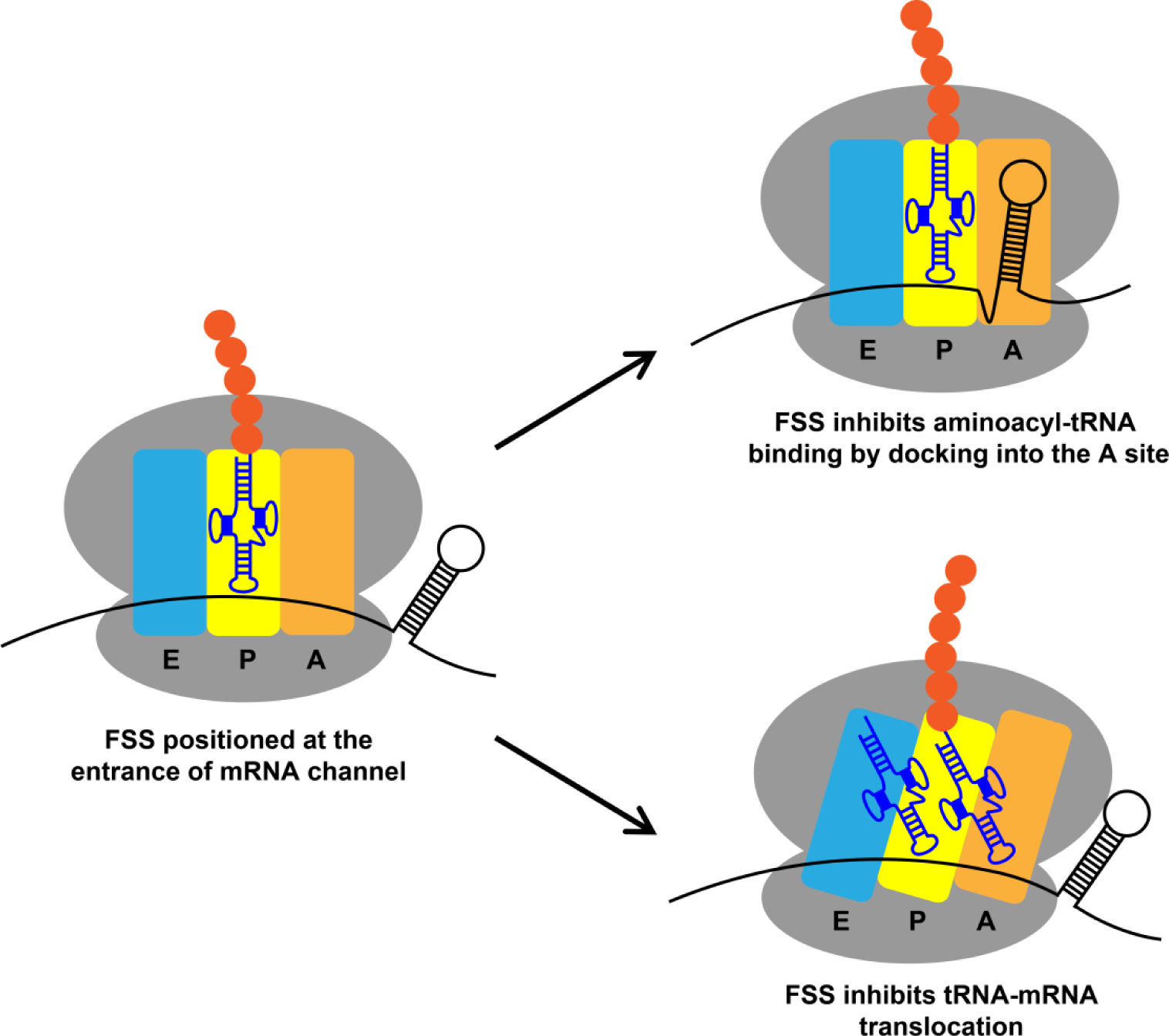
Two alternative mechanisms by which *dnaX* and HIV FSSs perturb translation elongation. Upon encountering the ribosome, the FSS can hinder tRNA binding by docking to the A site of the ribosome or inhibit translocation by interacting with the mRNA entry channel.

While dnaX and HIV FSSs dramatically perturbed the kinetics of the elongation cycle, our data provide no evidence that these stem-loops induce a unique conformation of the ribosome with a “super-rotated” orientation of ribosomal subunits reported previously (Qin et al., 2014). The super-rotated conformation, in which the ribosomal 30S subunit rotates by ∼20 degree against the 50S subunit, was inferred from smFRET data showing a 0.2 FRET value for the S6/L9 FRET pair when the ribosome encountered a dnaX FSS or mRNA/DNA duplex. In our smFRET study using ribosomes programmed with dnaX_Slip mRNA, we only detected fluctuations between 0.4 and 0.6 FRET states corresponding to R and NR conformations while no FRET states below 0.4 were observed. Previously observed 0.2 FRET of the S6/L9 FRET pair might correspond to nuclease-damaged ribosomes or be induced by the interaction of ribosomes with the microscope slide surface.

In our work, we observed 5-fold decrease of the rate of Lys-tRNA^Lys^ binding to the second Lys codon of dnaX slippery sequence induced by the FSS (Fig. 2). Furthermore, the rate of tRNA binding decreased by another order of magnitude when the slippery sequence of *dnaX* mRNA was replaced with non-slippery codons (Fig. 3-4). A-site tRNA inhibition was also observed with the HIV FSS in the context non-slippery codons (Fig. 3-4), suggesting a common underlying mechanism employed by a variety of FSSs. Such inhibition is only observed when the FSS is when positioned 11-12 nucleotides downstream of the P-site codon.

Our observation of FSS-dependent inhibition of A-site tRNA binding helps to resolve some inconsistencies in previous studies. One previous smFRET study suggested that the rate of A-site tRNA delivery during decoding of the slippery sequence of *dnaX* mRNA is unaffected by the presence of downstream FSS (Kim et al., 2014). Other single-molecule experiments suggested that Lys-tRNA^Lys^ accommodation during decoding of the second Lys codon of dnaX slippery sequence is delayed (Chen et al., 2014). Two-fold inhibition of A-site codon decoding induced by the FSS was observed in another study when dnaX slippery sequence was replaced with non-slippery Lys codons AAGAAG (Caliskan et al., 2017). Differences in experimental conditions (EF-Tu and tRNA concentrations) as well as sequence variations of model dnaX mRNAs may underlie inconsistencies between earlier studies of the FSS effect on tRNA binding. In particular, in aforementioned studies, the wild-type dnaX linker sequence AGUGA between the slippery sequence and the FSS was replaced with either UUUGA (Kim et al., 2014), UUCUA (Caliskan et al., 2017) or AGUUC (Chen et al., 2014).

When positioned 11-12 nucleotides downstream of the P-site codon, FSSs likely inhibit A-site tRNA binding and ribosome translocation by sampling two alternative conformations on the ribosome (Fig. 7), consistent with our observation of conformational heterogeneity of the hairpin in the A site. In one FSS conformation, which was previously seen in cryo-EM reconstruction of dnaX-ribosome complex (Zhang et al., 2018), the FSS interacts with the mRNA entry channel. It has been recently demonstrated that upon encountering mRNA secondary structure the ribosome translocates through two alternative (fast and slow) pathways (Desai et al., 2019). The interactions of FSSs with the mRNA entry channel may increase the flux through the slow pathway and thus decrease the average rate of ribosome translocation (Desai et al., 2019). In another FSS conformation, which was visualized by our single-particle cryo-EM analysis, nucleotides between the P-site and the FSS disassociate from the mRNA channel, and the HIV FSS docks into the A site thus sterically hindering tRNA binding.

The linker sequence between the A-site codon and the FSS may facilitate binding of the FSS into the A site. Our structure suggests that the homo-purine sequence encompassing the second and third positions of the A-site codon and the linker form a purine stack. The formation of an A-form like single-stranded helix by the linker nucleotides may make the release of the mRNA nucleotides between P-site codon and FSS from the mRNA channel and docking of the FSS into the A site more thermodynamically favorable.

The HIV FSS is the largest but not the first hairpin observed in the ribosomal A site. A short four-basepair-long hairpin was observed in the A site of the ribosome bound to a short model mRNA (MF36) derived from phage T4 gene 32 mRNA (Yusupova et al., 2001). Crystallographic analyses suggested that the hairpin following the start codon may facilitate initiation of translation of gene 32 (Yusupova et al., 2001). Another short five-basepair-long hairpin of the bacteriophage T4 gene 60 has been visualized by cryo-EM in the A-site of the ribosome, which was stalled at the “take-off” mRNA site containing a stop codon (Agirrezabala et al., 2017). The hairpin prevents binding of the release factor 1 (RF1), thus inhibiting translation termination and inducing translational bypassing (Agirrezabala et al., 2017). Another short two-basepair-long hairpin was observed in the A site of the ribosome in which the P and A sites were occupied by an inhibitory codon pair CGA-CCG that is known to cause ribosome stalling (Gamble et al., 2016; Tesina et al., 2020). These observations suggest that occlusion of the A site by RNA stem-loops may be a common strategy shared by many regulatory stem-loops. Our findings of hairpin competition with tRNA expose a novel mechanism that retroviruses, including HIV, likely employ to regulate viral gene expression and expand viral proteome via mRNA frameshifting. Transient occlusion of the A site by a mRNA hairpin may also underlie programmed ribosome pausing/stalling events that trigger targeting nascent polypeptide chain to membrane (Young and Andrews, 1996) and No-Go mRNA decay (Doma and Parker, 2006).

## Materials and Methods

### Ribosome, EF-G, EF-Tu and tRNA preparation

tRNA^fMet^, tRNA^Met^, tRNA^Phe^, tRNA^Val^, tRNA^Tyr^, tRNA^Lys^, and tRNA^Glu^ (purchased from Chemical Block) were aminoacylated as previously described (Lancaster & Noller 2005; Moazed & Noller 1989). Tight couple 70S ribosomes used for biochemical experiments and ribosomal subunit used for cryo-EM sample assembly were purified from *E.coli* MRE600 stain as previously described (Ermolenko et al., 2007). S6-Cy5/L9-Cy3 ribosomes were prepared by partial reconstitution of ΔS6-30S and ΔL9-50S subunits with S6-41C-Cy5 and L11-11C-Cy3 as previously described (Ermolenko et al., 2007; Ling and Ermolenko, 2015). Histidine-tagged EF-G and EF-Tu were expressed and purified using previously established procedures (Ermolenko et al., 2007).

### Preparation of model mRNAs

Sequences encoding dnaX and HIV mRNAs were cloned by directional cloning downstream of T7 promoter in pSP64 plasmid vector (Promega Co). Model mRNAs (Suppl. Table 1) were generated by T7 polymerase-catalyzed run-off *in vitro* transcription and purified by denaturing PAGE. Prior to transcription, 3’ ends of the model mRNAs were defined by linearizing the corresponding DNA templates at specific restriction sites (Suppl. Table 1).

### smFRET measurements

smFRET measurements were done as previously described (Cornish et al., 2008; Ling and Ermolenko, 2015) with modifications. The quartz slides used for total internal reflection fluorescence (TIRF) microscopy were treated with dichlorodimethylsilane (DDS) (Hua et al. 2014). The DDS surface was coated with biotinylated BSA (bio-BSA). Uncoated areas were then passivated by 0.2% Tween-20 prepared in H50 buffer which contained 20 mM HEPES (pH 7.5) and 50 mM KCl. 30 μL 0.2 mg/mL neutravidin (dissolved in H50 buffer) was bound to the biotin-BSA. For each flow-through chamber, non-specific sample binding to the slide was checked in the absence of neutravidin. Ribosomal complexes were imaged in polyamine buffer (50 mM HEPES (pH7.5), 6 mM Mg^2+^, 6 mM β-mercaptoethanol, 150 mM NH_4_Cl, 0.1 mM spermine and 2 mM spermidine) with 0.8 mg/mL glucose oxidase, 0.625% glucose, 1.5 mM 6-hydroxy-2,5,7,8-tetramethylchromane-2-carboxylic (Trolox) and 0.4 μg/mL catalase. smFRET data were acquired with 100 ms time resolution.

IDL software (ITT) was used to extract florescence intensities of Cy3 donor (*I_D_*) and Cy5 acceptor (*I_A_*), from which apparent FRET efficiency (*E_FRET_*, hence referred as FRET) was calculated:

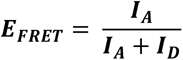

Traces showing single-step photobleachings for both Cy5 and Cy3 were selected using MATLAB scripts. FRET distribution histograms compiled from hundreds of smFRET traces were smoothed with a 5-point window using MATLAB and fit to two Gaussians corresponding to 0.4 and 0.6 FRET states (Ling & Ermolenko 2015; Cornish et al. 2008; Ermolenko et al. 2007). To determine rates of fluctuations between 0.4 and 0.6 FRET states, smFRET traces were idealized by 2-state Hidden Markov model (HMM) using HaMMy software and analyzed using transition density plot (TDP) analysis (McKinney et al. 2006).

Ribosome complexes used in smFRET experiments were assembled as follows. To fill the P site, 0.3 μM S6/L9-labeled ribosomes were incubated with 0.6 μM tRNA and 0.6 μM mRNA in polyamine buffer at 37 °C for 15 minutes. To bind aminoacyl-tRNA to the ribosomal A site, 0.6 μM aminoacyl-tRNA were pre-incubated with 10 μM EF-Tu and 1 mM GTP in polyamine buffer at 37 °C for 10 minutes. Then, 0.3 μM ribosomal complex containing peptidyl-tRNA in the P site was incubated with 0.6 μM aminoacyl-tRNA (complexed with EF-Tu•GTP) at 37 °C for 5 minutes. For the mixture of all *E.coli* tRNAs (Suppl. Fig. S), 30-(0.9 μM) or 150-fold (4.5 μM) molar excess of total aminoacyl-tRNAs (charged with all amino acids except for Tyr) were incubated with 30 nM ribosomes. After the incubation, ribosome samples were diluted to 1 nM with polyamine buffer, loaded on slide and immobilized by neutravidin and biotinylated DNA oligo annealed to the handle sequence of the ribosome-bound model mRNA. To catalyze translocation, 1 μM EF-G•GTP was added to the imaging buffer.

To prepare dnaX_Slip mRNA-programmed ribosomes that contained *N*-Ac-Val-Lys-tRNA^Lys^ in the P site (Fig. 2), *N*-Ac-Val-tRNA^Val^ and Lys-tRNA^Lys^ were bound to the P and A sites of the S6/L9-labeled ribosome, respectively, as described above. After complex immobilization on the slide and removal of unbound Lys-tRNA^Lys^, ribosomes were incubated with 1 μM EF-G•GTP at room temperature for 10 minutes. Next, EF-G•GTP was replaced with the imaging buffer and a mixture of 1 μM of EF-Tu•GTP•Lys-tRNALys and 1 μM EF-G•GTP (in imaging buffer) was delivered at 0.4 mL/min speed by a syringe pump (J-Kem Scientific) after 10 seconds of imaging.

### Puromycin assay

0.6 μM 70S ribosomes were incubated with 1.2 μM dnaX_NS mRNA and 1.2 μM *N*-Ac-[^3^H]-Phe-tRNA^Phe^ in polyamine buffer at 37 °C for 15 minutes followed by 10 minute incubation with 1 mM puromycin. The puromycin reaction was terminated by diluting the ribosome samples using MgSO_4_-saturated 0.3 M sodium acetate (pH 5.3), and the *N*-Ac-[^3^H]-Phe-puromycin was extracted ethyl acetate.

### Filter-binding assay

Filter-binding assay was performed as previously described (Salsi et al., 2016; Spiegel et al., 2007) with minor modifications. Ribosome complexes were assembled with radiolabeled tRNAs ([^14^C]Val-tRNA^Val^, [^3^H]Phe-tRNA^Phe^, [^3^H]Tyr-tRNA^Tyr^, [^3^H]Glu-tRNA^Glu^ and *N-*Ac-[^3^H]Tyr-tRNA^Tyr^ as indicated in figure legends) similarly to smFRET experiments described above. Ribosome complexes were applied to a nitrocellulose filter (MiliporeSigma), which was subsequently washed with 500 μl (for complexes programmed with dnaX mRNA) or 800 μl (for complexes programmed with HIV mRNA) of ice-cold polyamine buffer containing 20 mM Mg^2+^ to remove unbound tRNAs. 20 mM Mg^2+^ concentration was used to stabilize ribosome complexes under non-equilibrium conditions.

### HIV mRNA-70S ribosome complex assembly for cryo-EM analysis

The 70S ribosomes re-associated from 30S and 50S subunits were purified using sucrose gradient. 0.4 μM 70S ribosomes were bound with 0.7 μM *N*-Ac-Phe-tRNA^Phe^ and 0.8 μM HIV_NS (GAG) mRNA in polyamine buffer.

### Cryo-EM and Image Processing

C-flat grids (Copper, 1.2/1.3, Protochips) were glow-discharged for 30 sec in a PELCO glow-discharge unit at 15 mA. 3 μl of the 70S•HIV FSS-mRNA complex at 250 nM concentration were applied to the grid and incubated for 30 sec before vitrification using an FEI Vitrobot Mark IV (ThermoFisher). The grids were blotted for 3 sec using blotting force 3 at 4 °C and ∼90% humidity, plunged in liquid ethane, and stored in liquid nitrogen.

A dataset was collected using SerialEM (Mastronarde, 2005) on a Titan Krios operating at 300 kV and equipped with a K2 Summit camera (Gatan). A total of 5208 movies were collected using three shots per hole in super-resolution mode and a defocus range of −0.5 to −2.5 μm. The exposure length was 75 frames per movie yielding a total dose of 75 e-/ Å^2^. The super-resolution pixel size at the specimen level was 0.5115 Å. All movies were saved dark-corrected.

Gain and dark references were calculated using the method described by Afanasyev et al. (Afanasyev et al., 2015) and used to correct the collected movies in cisTEM (Grant et al., 2018). All further image processing was done using cisTEM. The movies were magnification-distortion-corrected using a calibrated distortion angle of 42.3° and a scale factor of 1.022 along the major axis and binned by a factor of 2. The movies were motion-corrected using all frames, and CTF parameters were estimated. Particles were picked using the particle picker tool in cisTEM and then curated manually. A total of 640,261 particles were extracted in 648^3^ pixel boxes.

Extracted particles were aligned to an unpublished reference volume using a global search in the resolution range from 8-300 Å (for classification workflow, see Supp. Fig. 5). The resulting 3D reconstruction was calculated using 50% of the particles with the highest scores and had a resolution of 3.27 Å (Fourier Shell Correlation FSC=0.143). Next, classification into 8 classes without alignment with a focus mask around the A-, and P-sites of the large and small subunit yielded two classes with density in the A-site. The classes corresponded to one rotated (23.15% of all particles), and one non-rotated state (11.44% of all particles), respectively. Both states were extracted separately and refined using local refinement with increasing resolution limits to 5 Å followed by one round of CTF refinement without alignment. The rotated and the non-rotated states reached resolutions of 3.15 Å and 3.35 Å, respectively. Each class then was classified into five classes without alignment using a focus mask around the observed density in the A-site. Two classes obtained from the non-rotated state showed weak density in the A-site. The two classes were merged and classified further into eight classes. Two classes had A-site density of which one showed strong density corresponding to the hairpin in the A-site and tRNA^Phe^ in the P-site. Particles for this class were extracted and aligned with increasing resolution limits to 5 Å. Finally, CTF refinement to 4 Å resolution without alignment and a step size of 50 Å was run and the final reconstructions were calculated using a beam-tilt corrected particle stack yielding final resolutions of 3.4 Å and 3.3 Å (Fourier Shell Correlation FSC=0.143).

The classification for the R conformation was done as described for the NR conformation. Classification into five classes yielded two classes with hairpin density. The classes were merged and classified into 8 classes of which four classes had weak density and one class yielded strong density. This class was extracted, CTF, and beam-tilt refined yielding a final resolution of 3.1 Å.

Finally, the obtained maps were sharpened in cisTEM and using the local resolution dependent function in phenix.autosharpen (Terwilliger et al., 2018).

### Model Building and Refinement

As the starting model for refinement we used the structure of the *E.coli* 70S ribosome with a ternary complex (PDB ID 5UYL), omitting EF-Tu and the A-site tRNA. An NMR structure of the HIV-1 frameshifting element (PDB ID 1PJY) was used as the starting model for the hairpin and to generate secondary-structure restraints. Missing parts of the mRNA were built manually and the geometry was regularized in phenix.geometry_minimization before refinement. The A-site finger was modelled using nucleotides 873-904 from PDB ID 5KPS where the A-site finger is well-ordered. Protein secondary structure restraints were generated in Phenix (Adams et al., 2010) and edited manually. We generated base-pairing (hydrogen bonds) using the “PDB to 3D Restraints” web-server (http://rna.ucsc.edu/pdbrestraints/, (Laurberg et al., 2008)) and added stacking restraints manually for the hairpin, and A-site finger.

Initially, the ribosomal subunits, tRNA and the hairpin were separately fitted into the cryo-EM, using Chimera, followed by manual adjustments in Coot (version 0.9-pre) (Emsley et al., 2010). The structural model was refined using phenix.real_space_refine (Afonine et al., 2018) and alternated with manual adjustments in Coot. The final model was evaluated in MolProbity (Williams et al., 2018).

## Supplementary Materials

### Ribosome stalling in the NR conformation is not due to frameshifting

Although the FSS alone is not expected to induce significant levels of frameshifting in the absence of a slippery sequence, we tested whether the stalling in the NR conformation observed with dnaX_NS and HIV_NS mRNAs was due to mRNA frameshifting that prevented A-site binding of Tyr-tRNA^Tyr^ (or Glu-tRNA^Glu^ in the case of ribosomes programmed with HIV_NS (GAG) mRNA). S6-cy5/L9-cy3 ribosomes, which were programmed with either dnaX_NS (UAC) or HIV_NS (UAC) mRNA and bound with P-site *N*-Ac-Phe-tRNA^Phe^ (Suppl. Fig. S4A, C), were incubated for 5 minutes with EF-Tu•GTP and 150-fold molar excess of total tRNA from *E.coli* aminoacylated with 19 natural amino acids except for Tyr. Incubation with total aa-tRNA (minus Tyr) did not lead to an appreciable increase in the fraction of the R (0.4 FRET) conformation (Suppl. Fig. S4B, D), indicating the lack of A-site tRNA binding in the absence of Tyr-tRNA^Tyr^. By contrast, as a positive control, just a 30-fold molar excess of total aa-tRNA (minus Tyr) was sufficient to decode an in-frame Glu (GAG) codon in ribosomes programmed with HIV_NS (GAG) ΔFSS mRNA as evident from the conversion of the ribosome population from the NR to R conformation (Suppl. Fig. S4E-G). Therefore, in the absence of the slippery sequence, FSS-induced frameshifting is negligible and does not account for ribosome stalling in the NR conformation observed in the experiments with ribosomes programmed with dnaX_NS or HIV_NS mRNAs.

### dnaX and HIV FSSs inhibit tRNA binding to the A site of the ribosome

HIV and dnaX FSSs placed near the entry to the mRNA channel could stall the ribosome in the NR conformation by either (i) inhibiting A-site tRNA binding, (ii) blocking the peptidyltransfer reaction after the binding of A-site tRNA or (iii) stabilizing the pretranslocation ribosome in the NR conformation. Pretranslocation-like S6-cy5/L9-cy3 ribosomes containing deacylated P-site tRNA^Phe^ exhibited similar intersubunit dynamics regardless of whether they were programmed with dnaX_NS, dnaX_NS ΔFSS, HIV_NS or HIV_NS ΔFSS mRNAs. These complexes fluctuated between the R (0.4 FRET) and NR (0.6 FRET) states at rates of 0.2-0.3 sec^-1^ (0.4 FRET to 0.6 FRET) and 0.7-0.8 sec^-1^, (0.6 FRET to 0.4 FRET), respectively, and spent 80% of time in the R conformation (Suppl. Fig. S3C, F, I, L). Hence, neither dnaX FSS nor HIV FSS placed near the entry of the mRNA channel directly affect intersubunit dynamics.

dnaX and HIV FSSs also did not change the sensitivity of P-site *N*-Ac-Phe-tRNA^Phe^ toward the A-site aminoacyl-tRNA mimic, antibiotic puromycin (Suppl. Fig. S5A), indicating that the frameshifting-inducing stem-loops placed at the entry of the mRNA channel do not block the transpeptidase activity of the ribosome. Therefore, in our smFRET experiments, FSSs from dnaX and HIV likely stall the ribosome in the NR conformation by inhibiting A-site tRNA binding.

---

**Supplementary Figure 1.**
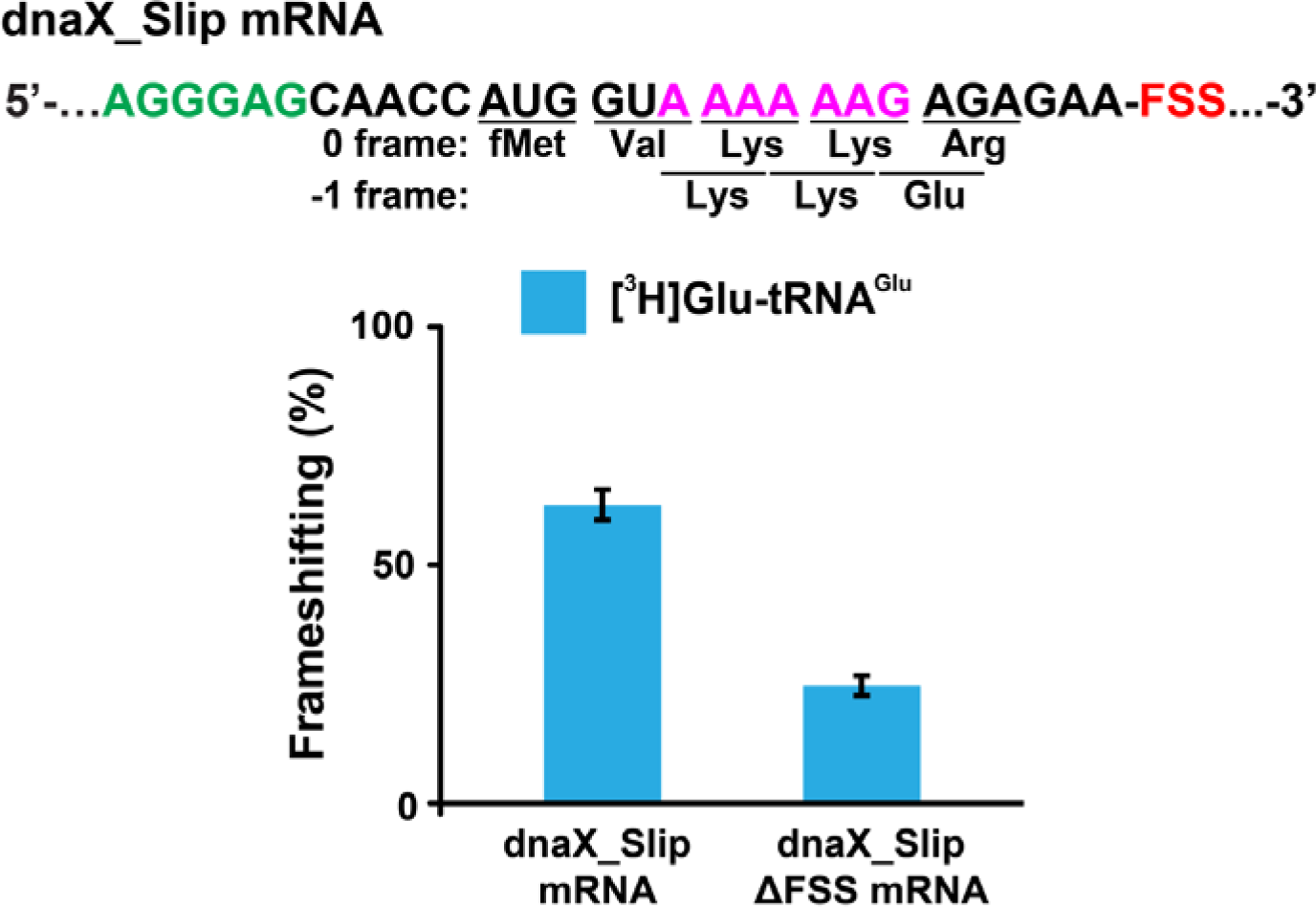
FSS and slippery sequence of *dnaX* mRNA stimulate −1 RPF. Ribosomes containing P-site *N*-Ac-Val-tRNA^Val^ were programmed with either dnaX_Slip or dnaX_Slip ΔFSS mRNA. The ribosomes were incubated with EF-G•GTP, EF-Tu•GTP, Lys-tRNA^Lys^, Arg-tRNA^Arg^ (binds in 0 frame) and [^3^H]Glu-tRNA^Glu^ (binds in - 1 frame) for 6 minutes. [^3^H]Glu-tRNA^Glu^ binding was measured by filter-binding assay. Frameshifting efficiency (ribosome occupancy by [^3^H]Glu-tRNA^Glu^) is shown relative to the P-site occupancy of *N*-Ac-[^3^H]Glu-tRNA^Glu^ in the ribosome programmed with dnaX_Slip ΔFSS mRNA. Error bars show standard deviations of triplicated measurements.

**Supplementary Figure 2.**
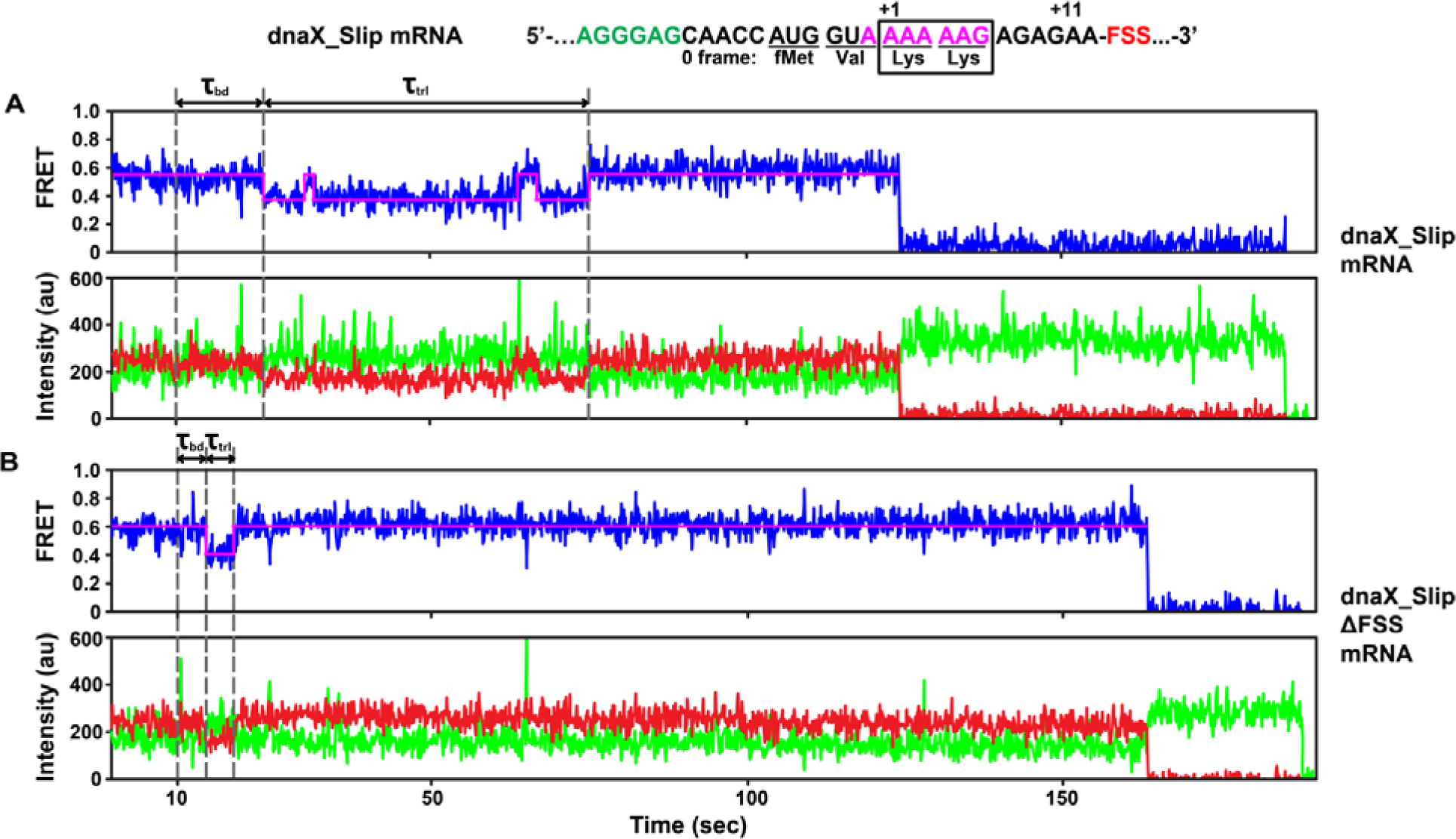
Both A-site tRNA binding and translocation are hindered by the dnaX FSS positioned at the entrance of mRNA 3wAchannel. (A-B) The full-length view of smFRET traces shown in Fig. 2 A-B with a single-step photobleaching of both Cy3 and Cy5 fluorophores.

**Supplementary Figure 3.**
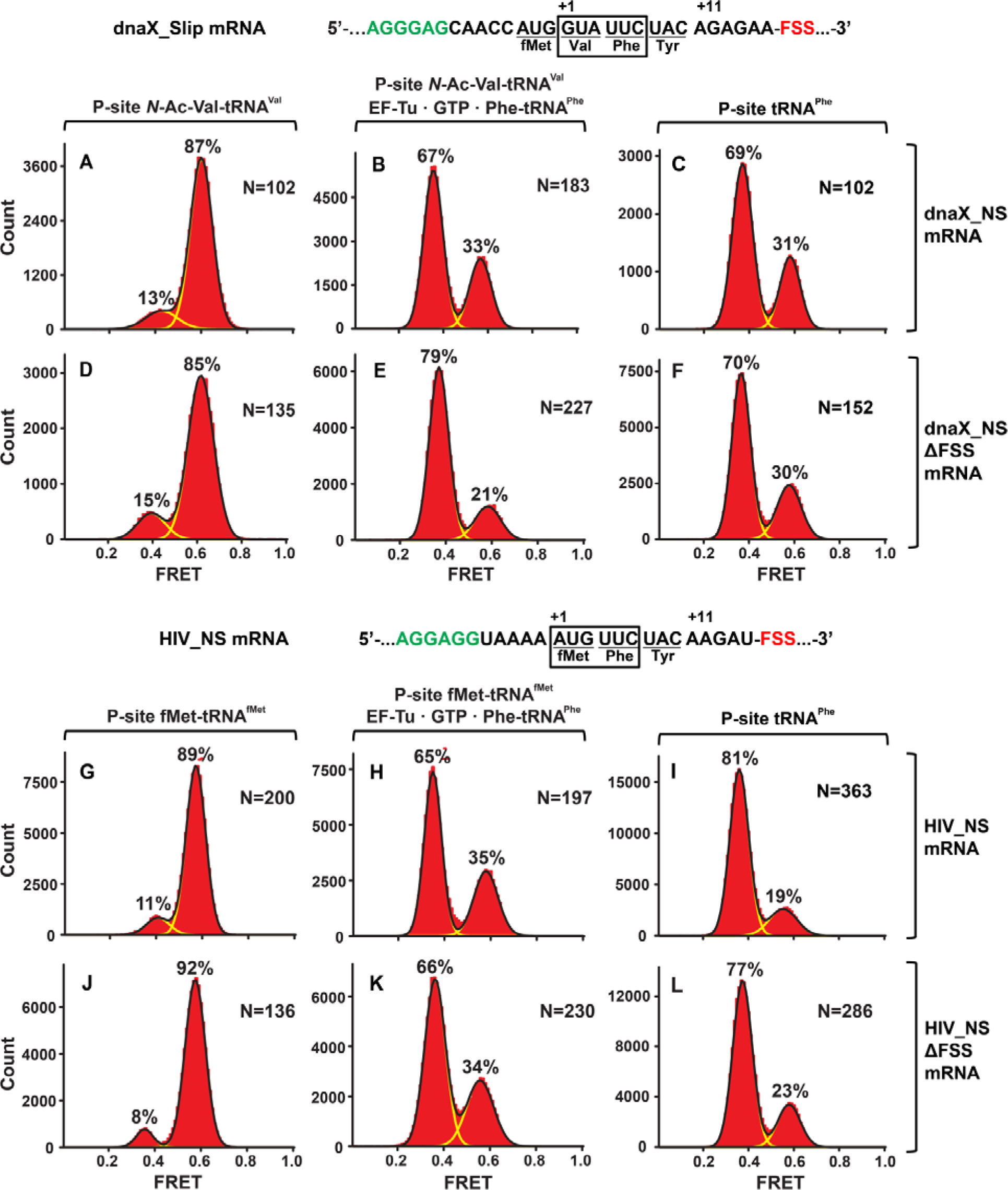
When positioned ≥3 nucleotides away from the entrance of mRNA channel at the beginning of elongation cycle, dnaX and HIV FSSs do not perturb ribosomal intersubunit rotations. Histograms show FRET distributions in S6-cy5/L9-cy3 ribosomes programmed with the dnaX_NS (A-C), dnaX_NS ΔFSS (D-F), HIV_NS (G-I), HIV_NS ΔFSS (J-L) mRNA, respectively. Ribosomes containing *N*-Ac-Val-tRNA^Val^ (A, D) or fMet-tRNA^fMet^ (G, J) in the P site were incubated with EF-Tu•GTP•Phe-tRNA^Phe^ for 5 minutes and imaged after removal of unbound Phe-tRNA^Phe^ (B, E, H, K). (C, F, I, L) Ribosomes containing deacylated tRNA^Phe^ in the P site.

**Supplementary Figure 4.**
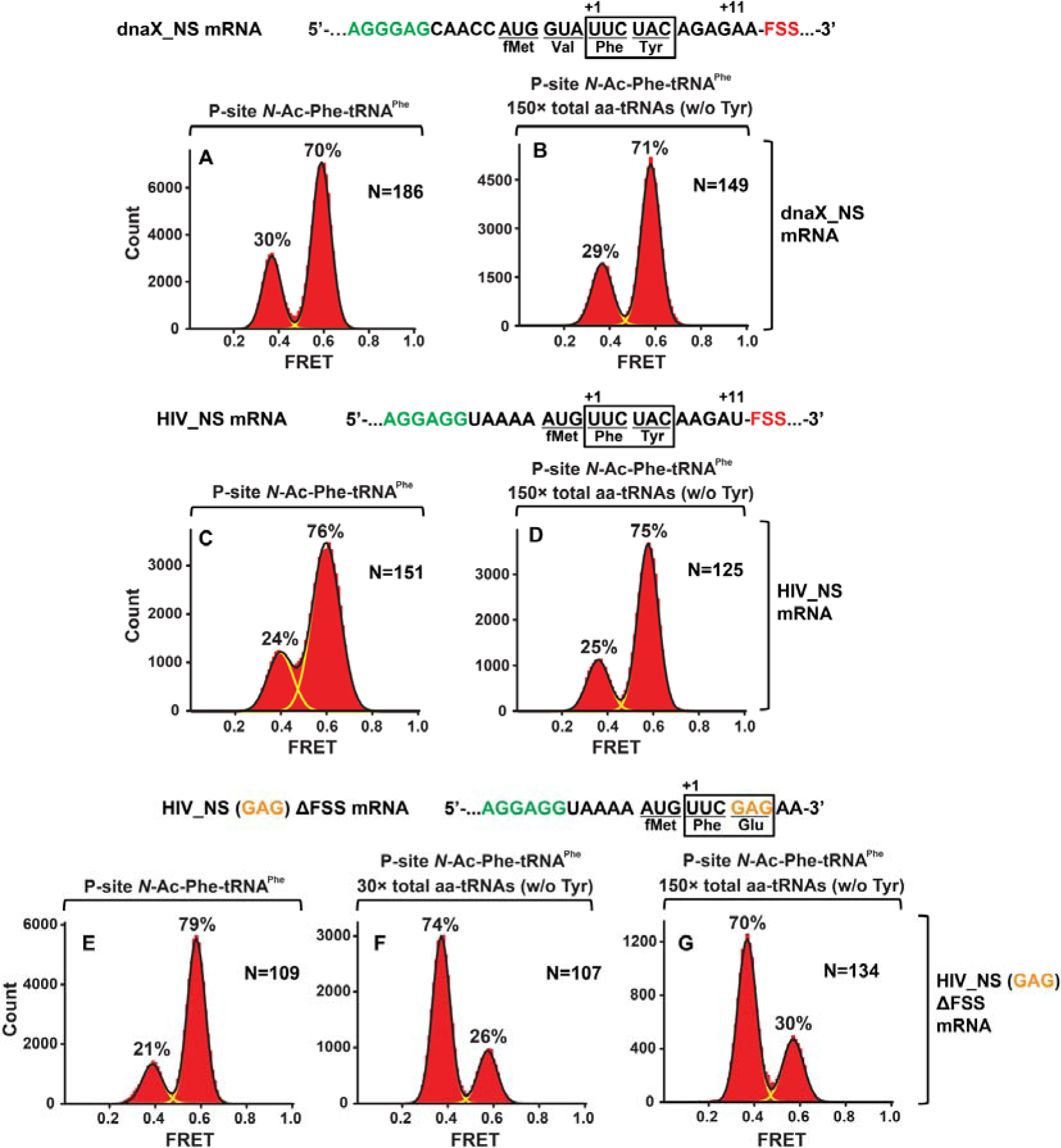
In the absence of slippery sequence, levels of the FSS-induced frameshifting are negligible. Histograms show FRET distributions in S6-cy5/L9-cy3 ribosomes programmed with the dnaX_NS (**A, B**), HIV_NS (**C, D**), or HIV_NS (GAG) ΔFSS (**E-G**) mRNA, respectively. Ribosomes containing P-site *N*-Ac-Phe-tRNA^Phe^ (**A, C, E**) were incubated with 150-(**B, D, G**) or 30-(**F**) fold molar excess of total aminoacyl-tRNAs•EF-Tu•GTP (mixture of all aminoacyl-tRNAs except Tyr-tRNA^Tyr^) for 5 minutes and imaged after removal of unbound aminoacyl-tRNA.

**Supplementary Figure 5.**
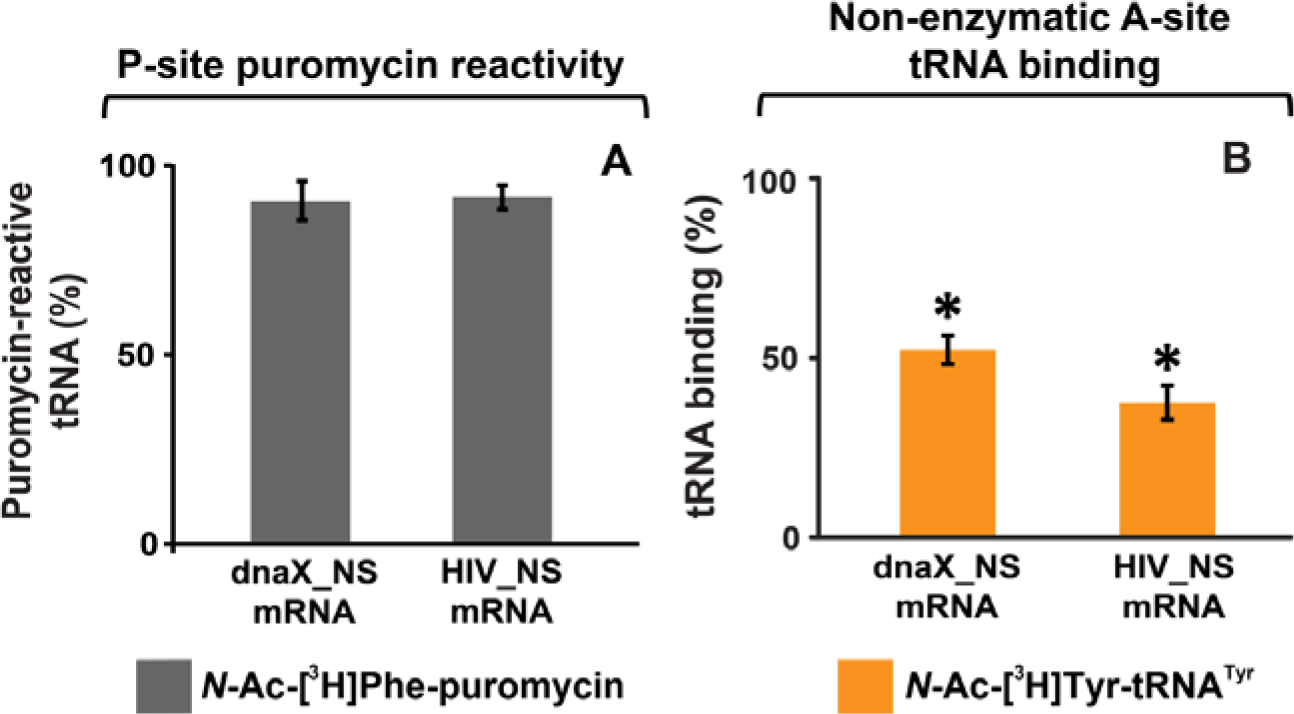
The FSS inhibits A-site tRNA binding but not ribosome-catalyzed transpeptidation reaction. (**A**) Ribosomes bound with P-site *N*-Ac-[^3^H]Phe-tRNA^Phe^ and FSS-containing mRNA were incubated with puromycin. The amount of *N*-Ac-[^3^H]Phe-puromycin extracted from the ribosomes programmed with dnaX_NS or HIV_NS mRNA was normalized by the amount of *N*-Ac-[^3^H]Phe-puromycin extracted from the ribosomes programmed with corresponding ΔFSS mRNAs. **(B)** The extent of *N*-Ac-[^3^H]Tyr-tRNA^Tyr^ binding to ribosomes bound with P-site deacylated tRNA^Phe^ and programmed with dnaX or HIV_NS mRNA after 10 minute incubation. The binding of *N*-Ac-[^3^H]Tyr-tRNA^Tyr^ determined by filter-binding assay is shown relative to that measured in ribosomes programmed with corresponding ΔFSS mRNAs. Asterisks indicate that amino acid incorporation into ribosomes programmed with FSS-containing mRNA was significantly different from that in ribosomes programmed with ΔFSS mRNA, as *p*-values determined by the Student t-test were below 0.05. Error bars in each panel show standard deviations of triplicated measurements.

**Supplementary Figure 6.**
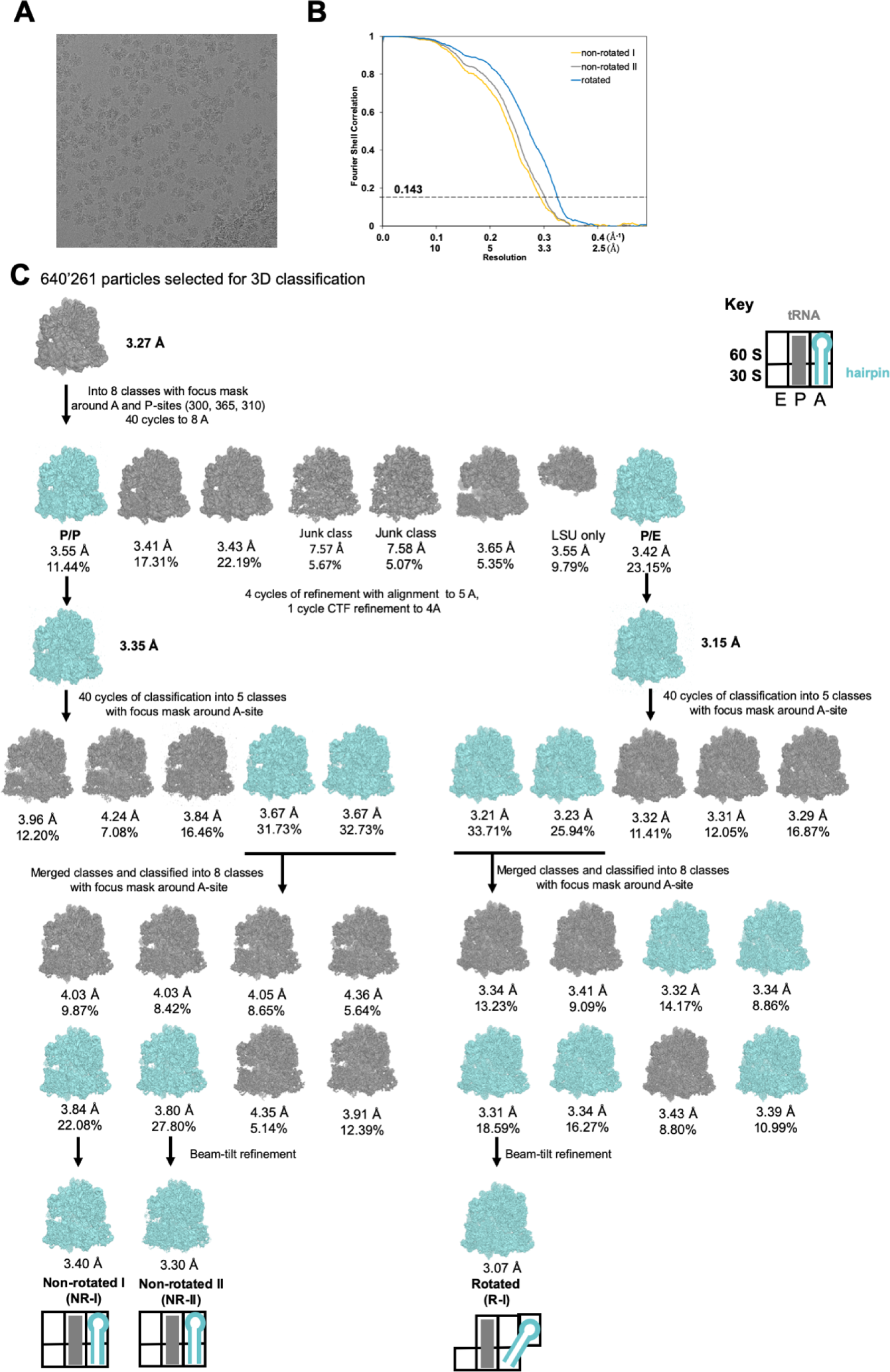
Data and schematic of cryo-EM refinement and classification. **(A)** Representative micrograph. **(B)** Fourier shell correlation as a function of resolution for the NR (I and II) and R structures. **(C)** 640,261 particles were aligned to a single model. Focused 3D classification using a spherical mask around the A and P sites yielded one class of R and one class of NR ribosomes. Each class was extracted and refined separately at 5 Å. Sub-classification of each class with a spherical mask around the A site yielded two classes of R and NR ribosomes with weak density for the HIV FSS hairpin in the A site. Particles were extracted based on HIV FSS hairpin occupancy and further sub-classified with a mask as described for the previous step.

**Supplementary Figure 7.**
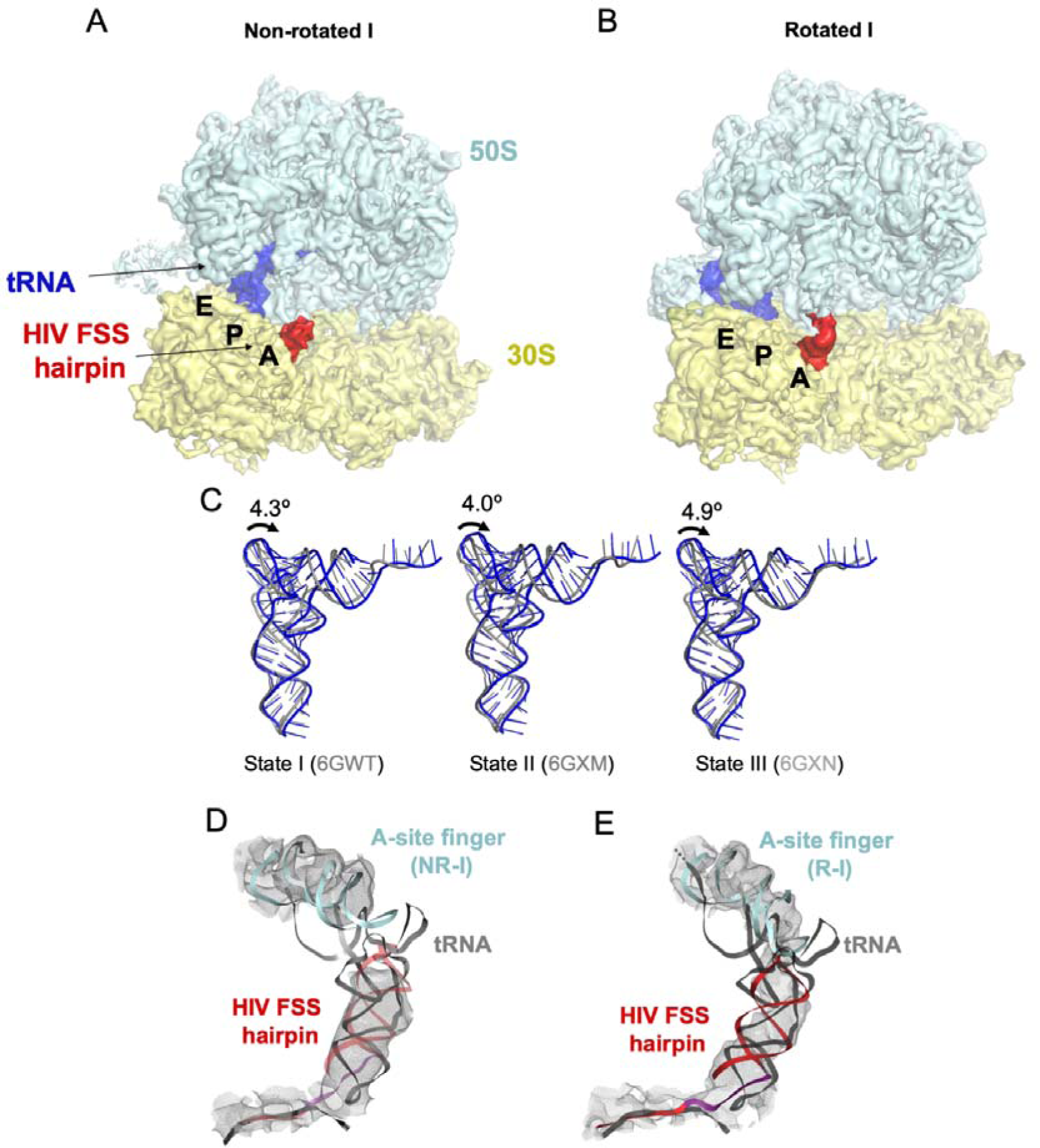
Density maps of **(A)** NR-I and **(B)** R-I filtered to 5 Å resolution. The cryo-EM maps are colored as in Figure 5. **(C)** Comparison of P-site tRNA of the NR-I structure from this work (blue) with tRNA from states I-III of recycling factor 1 and 3 (RF1 and RF3) bound ribosome structures stalled with Apidaecin 137 (grey) (Graf et al., 2018). **(D-E)** Fitting of a tRNA (grey) into the hairpin density of the (C) NR-I and (D) R-I states shows that tRNA cannot be accommodated without steric clashes with the 50S A-site finger (aqua) in either of the observed states. This rules out that the density corresponds to a tRNA.

**Supplementary Table 1.**
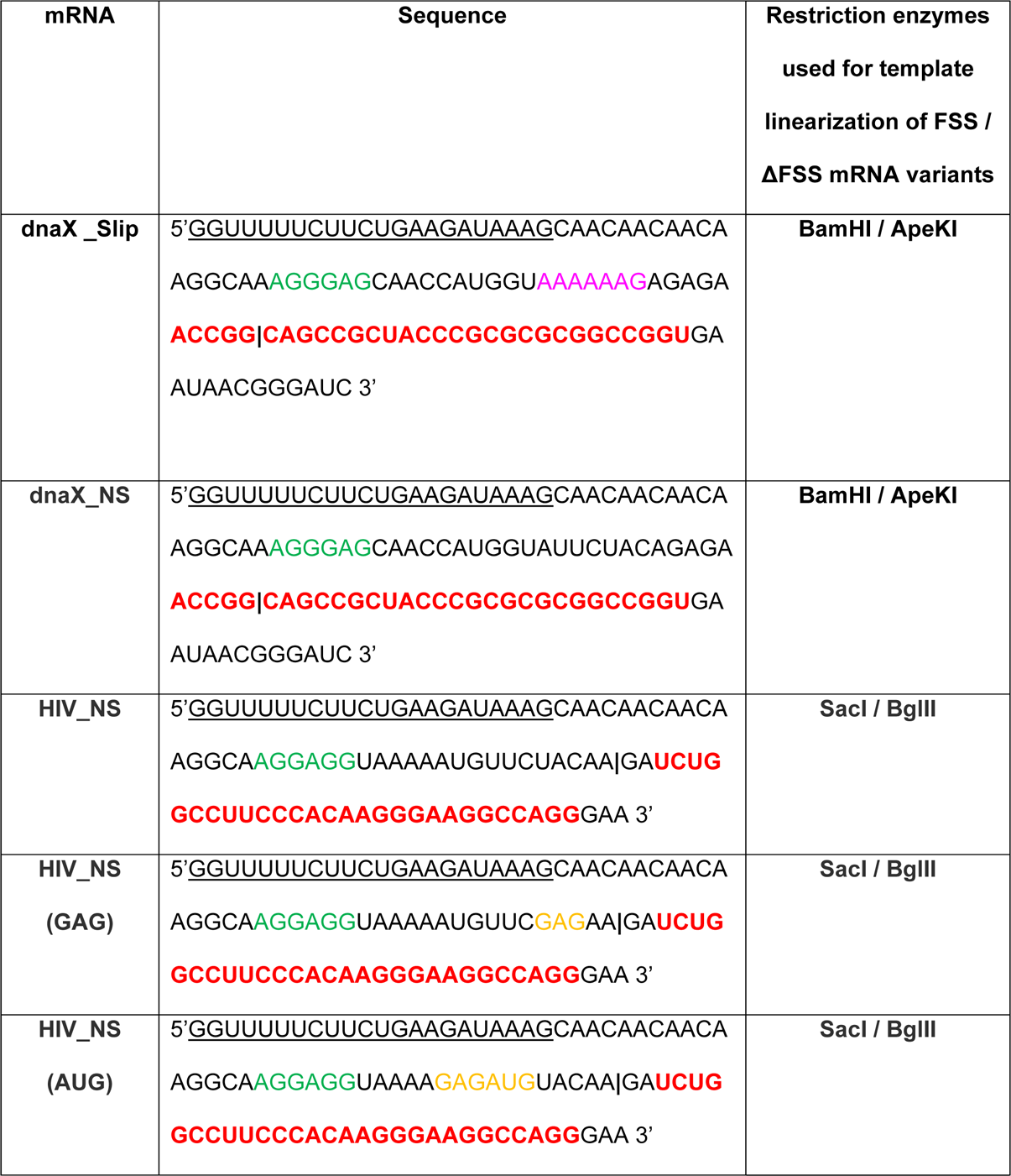
Model mRNA sequences. The SD sequence is shown in green, slippery sequence in magenta, FSS in red, and handle sequence, which is complementary to biotinylated DNA oligo, underlined. To alter codon identities, a UAC-to-GAG mutation was made on HIV_NS mRNA to generate HIV_NS (GAG) mRNA. Similarly, AUG-to-GAG UUC-to-AUG mutations were made to generate HIV_NS (AUG) mRNA. The codon replacements are colored orange. Vertical black bars indicate the 3’ ends of ΔFSS mRNAs.

## References

1. Adams, P.D., Afonine, P.V., Bunkoczi, G., Chen, V.B., Davis, I.W., Echols, N., Headd, J.J., Hung, L.W., Kapral, G.J., Grosse-Kunstleve, R.W., et al. (2010). PHENIX: a comprehensive Python-based system for macromolecular structure solution. Acta Crystallogr D Biol Crystallogr 66, 213–221.

2. Afanasyev, P., Ravelli, R.B., Matadeen, R., De Carlo, S., van Duinen, G., Alewijnse, B., Peters, P.J., Abrahams, J.P., Portugal, R.V., Schatz, M., et al. (2015). A posteriori correction of camera characteristics from large image data sets. Sci Rep 5, 10317.

3. Afonine, P.V., Poon, B.K., Read, R.J., Sobolev, O.V., Terwilliger, T.C., Urzhumtsev, A., and Adams, P.D. (2018). Real-space refinement in PHENIX for cryo-EM and crystallography. Acta Crystallogr D Struct Biol 74, 531–544.

4. Agirrezabala, X., Samatova, E., Klimova, M., Zamora, M., Gil-Carton, D., Rodnina, M.V., and Valle, M. (2017). Ribosome rearrangements at the onset of translational bypassing. Sci Adv 3, e1700147.

5. Aitken, C.E., and Puglisi, J.D. (2010). Following the intersubunit conformation of the ribosome during translation in real time. Nat Struct Mol Biol 17, 793–800.

6. Atkins, J.F., Baranov, P.V., Fayet, O., Herr, A.J., Howard, M.T., Ivanov, I.P., Matsufuji, S., Miller, W.A., Moore, B., Prere, M.F., et al. (2001). Overriding standard decoding: implications of recoding for ribosome function and enrichment of gene expression. Cold Spring Harb Symp Quant Biol 66, 217–232.

7. Belew, A.T., Meskauskas, A., Musalgaonkar, S., Advani, V.M., Sulima, S.O., Kasprzak, W.K., Shapiro, B.A., and Dinman, J.D. (2014). Ribosomal frameshifting in the CCR5 mRNA is regulated by miRNAs and the NMD pathway. Nature 512, 265–269.

8. Blanchard, S.C., Gonzalez, R.L., Kim, H.D., Chu, S., and Puglisi, J.D. (2004a). tRNA selection and kinetic proofreading in translation. Nat Struct Mol Biol 11, 1008–1014.

9. Blanchard, S.C., Kim, H.D., Gonzalez, R.L., Jr., Puglisi, J.D., and Chu, S. (2004b). tRNA dynamics on the ribosome during translation. Proc Natl Acad Sci U S A 101, 12893–12898.

10. Bock, L.V., Caliskan, N., Korniy, N., Peske, F., Rodnina, M.V., and Grubmuller, H. (2019). Thermodynamic control of −1 programmed ribosomal frameshifting. Nat Commun 10, 4598.

11. Brunelle, M.N., Payant, C., Lemay, G., and Brakier-Gingras, L. (1999). Expression of the human immunodeficiency virus frameshift signal in a bacterial cell-free system: influence of an interaction between the ribosome and a stem-loop structure downstream from the slippery site. Nucleic Acids Res 27, 4783–4791.

12. Caliskan, N., Katunin, V.I., Belardinelli, R., Peske, F., and Rodnina, M.V. (2014). Programmed −1 frameshifting by kinetic partitioning during impeded translocation. Cell 157, 1619–1631.

13. Caliskan, N., Peske, F., and Rodnina, M.V. (2015). Changed in translation: mRNA recoding by −1 programmed ribosomal frameshifting. Trends Biochem Sci 40, 265–274.

14. Caliskan, N., Wohlgemuth, I., Korniy, N., Pearson, M., Peske, F., and Rodnina, M.V. (2017). Conditional Switch between Frameshifting Regimes upon Translation of dnaX mRNA. Mol Cell 66, 558–567 e554.

15. Chen, C., Zhang, H., Broitman, S.L., Reiche, M., Farrell, I., Cooperman, B.S., and Goldman, Y.E. (2013). Dynamics of translation by single ribosomes through mRNA secondary structures. Nat Struct Mol Biol 20, 582–588.

16. Chen, G., Chang, K.Y., Chou, M.Y., Bustamante, C., and Tinoco, I., Jr. (2009). Triplex structures in an RNA pseudoknot enhance mechanical stability and increase efficiency of −1 ribosomal frameshifting. Proc Natl Acad Sci U S A 106, 12706–12711.

17. Chen, J., Petrov, A., Johansson, M., Tsai, A., O’Leary, S.E., and Puglisi, J.D. (2014). Dynamic pathways of −1 translational frameshifting. Nature 512, 328–332.

18. Choi, J., O’Loughlin, S., Atkins, J.F., and Puglisi, J.D. (2020). The energy landscape of −1 ribosomal frameshifting. Sci Adv 6, eaax6969.

19. Cornish, P.V., Ermolenko, D.N., Noller, H.F., and Ha, T. (2008). Spontaneous intersubunit rotation in single ribosomes. Mol Cell 30, 578–588.

20. Del Campo, C., Bartholomaus, A., Fedyunin, I., and Ignatova, Z. (2015). Secondary Structure across the Bacterial Transcriptome Reveals Versatile Roles in mRNA Regulation and Function. PLoS Genet 11, e1005613.

21. Desai, V.P., Frank, F., Lee, A., Righini, M., Lancaster, L., Noller, H.F., Tinoco, I., Jr., and Bustamante, C. (2019). Co-temporal Force and Fluorescence Measurements Reveal a Ribosomal Gear Shift Mechanism of Translation Regulation by Structured mRNAs. Mol Cell 75, 1007–1019 e1005.

22. Doma, M.K., and Parker, R. (2006). Endonucleolytic cleavage of eukaryotic mRNAs with stalls in translation elongation. Nature 440, 561–564.

23. Dunkle, J.A., Wang, L., Feldman, M.B., Pulk, A., Chen, V.B., Kapral, G.J., Noeske, J., Richardson, J.S., Blanchard, S.C., and Cate, J.H. (2011). Structures of the bacterial ribosome in classical and hybrid states of tRNA binding. Science 332, 981–984.

24. Emsley, P., Lohkamp, B., Scott, W.G., and Cowtan, K. (2010). Features and development of Coot. Acta Crystallogr D Biol Crystallogr 66, 486–501.

25. Ermolenko, D.N., Majumdar, Z.K., Hickerson, R.P., Spiegel, P.C., Clegg, R.M., and Noller, H.F. (2007). Observation of intersubunit movement of the ribosome in solution using FRET. J Mol Biol 370, 530–540.

26. Ermolenko, D.N., and Noller, H.F. (2011). mRNA translocation occurs during the second step of ribosomal intersubunit rotation. Nat Struct Mol Biol 18, 457–462.

27. Frank, J., and Agrawal, R.K. (2000). A ratchet-like inter-subunit reorganization of the ribosome during translocation. Nature 406, 318–322.

28. Frank, J., and Gonzalez, R.L., Jr. (2010). Structure and dynamics of a processive Brownian motor: the translating ribosome. Annu Rev Biochem 79, 381–412.

29. Gamble, C.E., Brule, C.E., Dean, K.M., Fields, S., and Grayhack, E.J. (2016). Adjacent Codons Act in Concert to Modulate Translation Efficiency in Yeast. Cell 166, 679–690.

30. Graf, M., Huter, P., Maracci, C., Peterek, M., Rodnina, M.V., and Wilson, D.N. (2018). Visualization of translation termination intermediates trapped by the Apidaecin 137 peptide during RF3-mediated recycling of RF1. Nat Commun 9, 3053.

31. Grant, T., Rohou, A., and Grigorieff, N. (2018). cisTEM, user-friendly software for single-particle image processing. Elife 7.

32. Hansen, T.M., Reihani, S.N., Oddershede, L.B., and Sorensen, M.A. (2007). Correlation between mechanical strength of messenger RNA pseudoknots and ribosomal frameshifting. Proc Natl Acad Sci U S A 104, 5830–5835.

33. Isaksson, J., Acharya, S., Barman, J., Cheruku, P., and Chattopadhyaya, J. (2004). Single-stranded adenine-rich DNA and RNA retain structural characteristics of their respective double-stranded conformations and show directional differences in stacking pattern. Biochemistry 43, 15996–16010.

34. Jacks, T., Power, M.D., Masiarz, F.R., Luciw, P.A., Barr, P.J., and Varmus, H.E. (1988). Characterization of ribosomal frameshifting in HIV-1 gag-pol expression. Nature 331, 280–283.

35. Johansson, M., Bouakaz, E., Lovmar, M., and Ehrenberg, M. (2008). The kinetics of ribosomal peptidyl transfer revisited. Mol Cell 30, 589–598.

36. Juette, M.F., Terry, D.S., Wasserman, M.R., Altman, R.B., Zhou, Z., Zhao, H., and Blanchard, S.C. (2016). Single-molecule imaging of non-equilibrium molecular ensembles on the millisecond timescale. Nat Methods 13, 341–344.

37. Kim, H.K., Liu, F., Fei, J., Bustamante, C., Gonzalez, R.L., Jr., and Tinoco, I., Jr. (2014). A frameshifting stimulatory stem loop destabilizes the hybrid state and impedes ribosomal translocation. Proc Natl Acad Sci U S A 111, 5538–5543.

38. Kim, H.K., and Tinoco, I., Jr. (2017). EF-G catalyzed translocation dynamics in the presence of ribosomal frameshifting stimulatory signals. Nucleic Acids Res 45, 2865–2874.

39. Kontos, H., Napthine, S., and Brierley, I. (2001). Ribosomal pausing at a frameshifter RNA pseudoknot is sensitive to reading phase but shows little correlation with frameshift efficiency. Mol Cell Biol 21, 8657–8670.

40. Korniy, N., Goyal, A., Hoffmann, M., Samatova, E., Peske, F., Pohlmann, S., and Rodnina, M.V. (2019). Modulation of HIV-1 Gag/Gag-Pol frameshifting by tRNA abundance. Nucleic Acids Res 47, 5210–5222.

41. Larsen, B., Gesteland, R.F., and Atkins, J.F. (1997). Structural probing and mutagenic analysis of the stem-loop required for Escherichia coli dnaX ribosomal frameshifting: programmed efficiency of 50%. J Mol Biol 271, 47–60.

42. Laurberg, M., Asahara, H., Korostelev, A., Zhu, J., Trakhanov, S., and Noller, H.F. (2008). Structural basis for translation termination on the 70S ribosome. Nature 454, 852–857.

43. Leger, M., Sidani, S., and Brakier-Gingras, L. (2004). A reassessment of the response of the bacterial ribosome to the frameshift stimulatory signal of the human immunodeficiency virus type 1. RNA 10, 1225–1235.

44. Ling, C., and Ermolenko, D.N. (2015). Initiation factor 2 stabilizes the ribosome in a semirotated conformation. Proc Natl Acad Sci U S A 112, 15874–15879.

45. Lopinski, J.D., Dinman, J.D., and Bruenn, J.A. (2000). Kinetics of ribosomal pausing during programmed −1 translational frameshifting. Mol Cell Biol 20, 1095–1103.

46. Mastronarde, D.N. (2005). Automated electron microscope tomography using robust prediction of specimen movements. J Struct Biol 152, 36–51.

47. Mazauric, M.H., Seol, Y., Yoshizawa, S., Visscher, K., and Fourmy, D. (2009). Interaction of the HIV-1 frameshift signal with the ribosome. Nucleic Acids Res 37, 7654–7664.

48. Moazed, D., and Noller, H.F. (1989). Intermediate states in the movement of transfer RNA in the ribosome. Nature 342, 142–148.

49. Mouzakis, K.D., Lang, A.L., Vander Meulen, K.A., Easterday, P.D., and Butcher, S.E. (2013). HIV-1 frameshift efficiency is primarily determined by the stability of base pairs positioned at the mRNA entrance channel of the ribosome. Nucleic Acids Res 41, 1901–1913.

50. Polikanov, Y.S., Steitz, T.A., and Innis, C.A. (2014). A proton wire to couple aminoacyl-tRNA accommodation and peptide-bond formation on the ribosome. Nat Struct Mol Biol 21, 787–793.

51. Qin, P., Yu, D., Zuo, X., and Cornish, P.V. (2014). Structured mRNA induces the ribosome into a hyper-rotated state. EMBO Rep 15, 185–190.

52. Qu, X., Wen, J.D., Lancaster, L., Noller, H.F., Bustamante, C., and Tinoco, I., Jr. (2011). The ribosome uses two active mechanisms to unwind messenger RNA during translation. Nature 475, 118–121.

53. Ritchie, D.B., Foster, D.A., and Woodside, M.T. (2012). Programmed −1 frameshifting efficiency correlates with RNA pseudoknot conformational plasticity, not resistance to mechanical unfolding. Proc Natl Acad Sci U S A 109, 16167–16172.

54. Rodnina, M.V., and Wintermeyer, W. (2001). Fidelity of aminoacyl-tRNA selection on the ribosome: kinetic and structural mechanisms. Annu Rev Biochem 70, 415–435.

55. Salsi, E., Farah, E., and Ermolenko, D.N. (2016). EF-G Activation by Phosphate Analogs. J Mol Biol 428, 2248–2258.

56. Sharma, H., Adio, S., Senyushkina, T., Belardinelli, R., Peske, F., and Rodnina, M.V. (2016). Kinetics of Spontaneous and EF-G-Accelerated Rotation of Ribosomal Subunits. Cell Rep 16, 2187–2196.

57. Somogyi, P., Jenner, A.J., Brierley, I., and Inglis, S.C. (1993). Ribosomal pausing during translation of an RNA pseudoknot. Mol Cell Biol 13, 6931–6940.

58. Spiegel, P.C., Ermolenko, D.N., and Noller, H.F. (2007). Elongation factor G stabilizes the hybrid-state conformation of the 70S ribosome. RNA 13, 1473–1482.

59. Staple, D.W., and Butcher, S.E. (2003). Solution structure of the HIV-1 frameshift inducing stem-loop RNA. Nucleic Acids Res 31, 4326–4331.

60. Svidritskiy, E., Demo, G., Loveland, A.B., Xu, C., and Korostelev, A.A. (2019). Extensive ribosome and RF2 rearrangements during translation termination. Elife 8.

61. Takyar, S., Hickerson, R.P., and Noller, H.F. (2005). mRNA helicase activity of the ribosome. Cell 120, 49–58.

62. Terwilliger, T.C., Sobolev, O.V., Afonine, P.V., and Adams, P.D. (2018). Automated map sharpening by maximization of detail and connectivity. Acta Crystallogr D Struct Biol 74, 545–559.

63. Tesina, P., Lessen, L.N., Buschauer, R., Cheng, J., Wu, C.C., Berninghausen, O., Buskirk, A.R., Becker, T., Beckmann, R., and Green, R. (2020). Molecular mechanism of translational stalling by inhibitory codon combinations and poly(A) tracts. EMBO J 39, e103365.

64. Tsuchihashi, Z., and Brown, P.O. (1992). Sequence requirements for efficient translational frameshifting in the Escherichia coli dnaX gene and the role of an unstable interaction between tRNA(Lys) and an AAG lysine codon. Genes Dev 6, 511–519.

65. Tsuchihashi, Z., and Kornberg, A. (1990). Translational frameshifting generates the gamma subunit of DNA polymerase III holoenzyme. Proc Natl Acad Sci U S A 87, 2516–2520.

66. Tu, C., Tzeng, T.H., and Bruenn, J.A. (1992). Ribosomal movement impeded at a pseudoknot required for frameshifting. Proc Natl Acad Sci U S A 89, 8636–8640.

67. Valle, M., Zavialov, A., Sengupta, J., Rawat, U., Ehrenberg, M., and Frank, J. (2003). Locking and unlocking of ribosomal motions. Cell 114, 123–134.

68. Wen, J.D., Lancaster, L., Hodges, C., Zeri, A.C., Yoshimura, S.H., Noller, H.F., Bustamante, C., and Tinoco, I. (2008). Following translation by single ribosomes one codon at a time. Nature 452, 598–603.

69. Williams, C.J., Headd, J.J., Moriarty, N.W., Prisant, M.G., Videau, L.L., Deis, L.N., Verma, V., Keedy, D.A., Hintze, B.J., Chen, V.B., et al. (2018). MolProbity: More and better reference data for improved all-atom structure validation. Protein Sci 27, 293–315.

70. Yan, S., Wen, J.D., Bustamante, C., and Tinoco, I., Jr. (2015). Ribosome excursions during mRNA translocation mediate broad branching of frameshift pathways. Cell 160, 870–881.

71. Young, J.C., and Andrews, D.W. (1996). The signal recognition particle receptor alpha subunit assembles co-translationally on the endoplasmic reticulum membrane during an mRNA-encoded translation pause in vitro. EMBO J 15, 172–181.

72. Yusupova, G.Z., Yusupov, M.M., Cate, J.H., and Noller, H.F. (2001). The path of messenger RNA through the ribosome. Cell 106, 233–241.

73. Zhang, Y., Hong, S., Ruangprasert, A., Skiniotis, G., and Dunham, C.M. (2018). Alternative Mode of E-Site tRNA Binding in the Presence of a Downstream mRNA Stem Loop at the Entrance Channel. Structure 26, 437-445 e433.

74. Zuker, M. (2003). Mfold web server for nucleic acid folding and hybridization prediction. Nucleic Acids Res 31, 3406–3415.

